# Dopaminergic processes predict temporal distortions in event memory

**DOI:** 10.1101/2025.05.14.654133

**Authors:** Erin Morrow, Ringo Huang, David Clewett

**Affiliations:** Department of Psychology, University of California, Los Angeles, USA 90095

**Keywords:** episodic memory, time, context, blinks, dopamine, event boundary

## Abstract

Our memories do not simply keep time — they warp it, bending the past to fit the structure of our experiences. For example, people tend to remember items as occurring farther apart in time if they spanned a change in context, or ‘event boundary,’ compared to the same context. While these distortions could sacrifice precise timing, they might also serve to divide and organize information into distinct memories. In the current study, we combined functional magnetic resonance imaging (fMRI; *n* = 32) with eye-tracking (*n* = 28) to test whether activation of the dopaminergic system, known to influence encoding and time perception, predicts time dilation between adjacent events in memory. Participants encoded item sequences while listening to tones that mostly repeated over time, forming a stable auditory context, but occasionally switched, creating an event boundary. We found that boundaries predicted greater retrospective estimates of time between item pairs. Critically, tone switches significantly activated the ventral tegmental area (VTA), a key midbrain dopaminergic region, and these responses predicted greater time dilation between item pairs that spanned those switches. Boundaries furthermore predicted a momentary increase in blinks. Activation of the VTA predicted blinking in general, consistent with the idea that blink behavior is a potential marker of dopaminergic activity. On a larger timescale, higher blink counts predicted greater time dilation in memory, but only for boundary-spanning item pairs. Together, these findings suggest that dopaminergic processes are sensitive to event structure and may drive temporal distortions that help to separate memories of distinct events.

## Introduction

“[Mechanical time] is unyielding, predetermined. [Body time] makes up its mind as it goes along,” remarks physicist and author Alan Lightman (1992, p.32). Indeed, time can seem malleable when we reflect upon the past. For example, experiences from the COVID-19 pandemic are not equally represented in memory. The early lockdown period is often recalled as having passed more quickly than it was originally experienced, potentially driven by a significant lack of variety or novelty (Droit-Volet et al., 2023; also see Rouhani et al., 2023 for similar findings). In contrast, everyday life typically involves frequent shifts in context, or ‘event boundaries’, which tend to stretch remembered time and promote the separation of experiences into distinct memories. These memory-expanding effects have been reliably demonstrated in laboratory settings, with changes in scenes (Ezzyat & Davachi, 2014), goals (Cowan et al., 2024), and emotional states (McClay, Sachs, & Clewett, 2023; Wang & Lapate, 2024) eliciting time dilation in long-term memory. Yet, despite their central role in human cognition, it remains unclear how the brain uses contextual cues to reshape temporal representations in service of memory organization.

A promising candidate for this process is the ventral tegmental area (VTA), a brainstem nucleus that releases dopamine and modulates episodic memory. Dopamine facilitates encoding and consolidation of behaviorally relevant information over both short (e.g., D’Ardenne et al., 2012) and long timescales (see Clewett & Murty, 2019; Harley, 2004). The VTA responds to a wide variety of salient stimuli, including those that are novel (Duszkiewicz et al., 2019) and carry predictive salience (Shohamy & Adcock, 2010). Neuroimaging studies in humans suggest that phasic VTA activation and dopamine may signal key moments when ongoing predictions fail and it is beneficial to update one’s mental model of what is happening (Antony et al., 2021; Zacks et al., 2011). Indeed, this framework is consistent with dopamine’s known role in working memory, which includes modulation of striatal activity to update ongoing representations (see D’Esposito & Postle, 2015) held in short-term memory storage (Baddeley & Hitch, 1974). Additionally, midbrain dopamine regulates synaptic plasticity in the hippocampus (Lisman & Grace, 2005; Shohamy & Adcock, 2010), a region that triggers the encoding of new memories and their temporal features (see e.g., Howard & Eichenbaum, 2015). Together, these factors make the VTA well equipped to mediate the influence of event boundaries on the temporal structure of memory.

Alongside its role in memory, dopamine is heavily implicated in time perception. According to the dopamine clock hypothesis (Meck, 1983; 1996), transient increases in dopamine may accelerate a hypothetical internal clock that generates pulses, or ‘ticks’, that are collected and stored by a memory system to estimate time. The more pulses collected during a fixed interval, the longer that period is perceived to last (Treisman, 1963). Thus, dopamine-induced acceleration of this clock would result in time overestimation (see Fung, Sutlief, & Hussain Shuler, 2022). Other frameworks offer additional nuance; for example, dopamine may modulate this clock based on prediction errors (Mikhael & Gershman, 2019), which are theorized to trigger event segmentation (Zacks et al., 2007) (see Fung, Sutlief, & Hussain Shuler, 2022). Many timing models also contain a working memory storage component that affects time perception (Fung, Sutlief, & Hussain Shuler, 2022) and is influenced by dopaminergic processes. However, most research has focused on how dopamine influences time perception as an experience unfolds, using tasks in which participants actively track the duration of an ongoing stimulus or estimate duration immediately after the stimulus ends. Therefore, it remains unclear whether dopamine also helps to encode exaggerated representations of time into long-term memory.

Intriguingly, blink behavior may offer an indirect window into the influence of dopamine and event boundaries on temporal memory distortions. Recent neuroimaging work shows that momentary blinks correspond with increased activation in dopaminergic regions, including the VTA and substantia nigra (Demiral et al., 2023). Moreover, it has been shown that spontaneous blink rate is altered in individuals with dopamine-related neuropsychiatric conditions like Parkinson’s disease (Fitzpatrick et al., 2012) and schizophrenia (Chen et al., 1996) under certain conditions (for review, see Jongkees & Colzato, 2016; Karson, 1983). Like dopamine, blinks also tend to occur around ‘breakpoints’ in experience, such as punctuation marks in text passages (Hall, 1945), pauses in speech (Nakano & Kitazawa, 2010), and the beginnings of quiz questions on a mentally demanding TV game show (Wyly et al., 2023) (also see Nakano et al., 2009; Nakano et al., 2013). Additionally, one study in humans showed that increased blinking during a short interval was associated with exaggerated estimates of time (Terhune, Sullivan, & Simola, 2016), consistent with the role of dopamine in shaping time perception.

In the current neuroimaging and eye-tracking study, we find that context shifts during item sequences predict increases in two indirect measures of dopaminergic processing: VTA activation and blinking. Irrespective of contextual stability or change, activation of the VTA is tightly coupled with momentary increases in blinking, supporting the putative relationship between dopamine and blink behavior. We also find that VTA activation and more prolonged periods of blinking across event boundaries predict the later expansion of temporal representations in memory. These findings extend the relationship between dopamine and time perception to episodic memory, revealing how the brain not only remembers time, but may also bend it to represent contextually distinct events.

## Results

### Event boundaries predicted subjective time dilation effects in long-term memory, a behavioral index of event separation

Participants performed a modified version of a paradigm known to induce temporal distortions in event memory (Clewett et al., 2020). Participants studied sequences of neutral object images, each preceded by a 1-s pure tone presented to their left or right ear (**Figure 1**). The laterality of the tone remained the same for eight items, creating the perception of a stable context or ‘event.’ However, after 8 successive items in each sequence, the tone abruptly switched to the other ear and changed pitch, forming a perceptual ‘event boundary.’ Participants were also instructed to switch the hand they used to respond to an orienting question for each image (“Is this object larger or smaller than a standard shoebox?”; for details on response distributions, see **Supplementary Figure 3**). In this way, event boundaries were not only perceptually salient but also task relevant. Each sequence contained three event boundaries, leading to four 8-item events.

**Figure 1.**
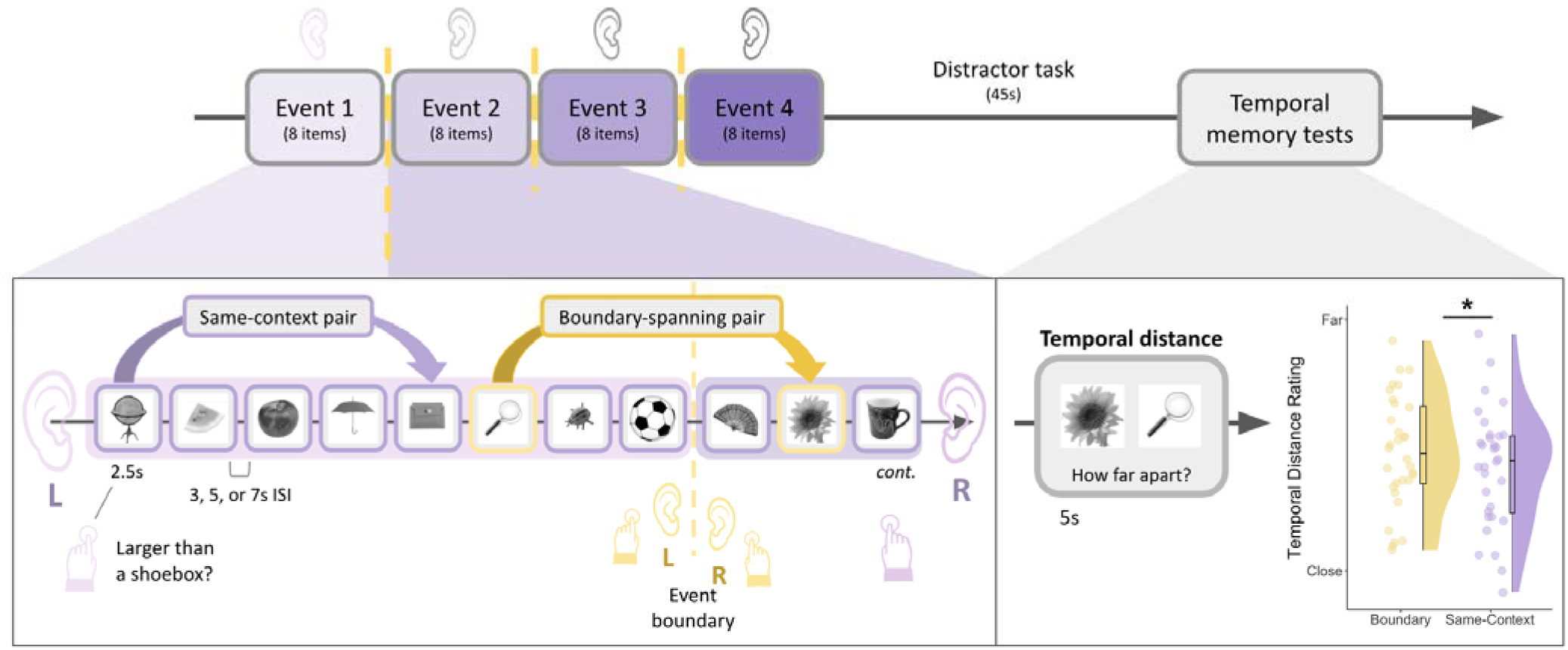
Event sequence encoding task and temporal distance memory test. In each block of the task, participants studied a sequence of 32 neutral object images. Prior to each image, participants listened to pure 1s tones either in their left or right ear during the midpoint of a fixation screen. The side of the tone also cued participants as to which hand they should use to judge whether each object was larger than a shoebox (e.g., left ear = left hand). Critically, the image sequence was divided into four events. Events were defined by a stable auditory context, with the same tone presented to the same ear for 8 successive items. After 8 items, the tone switched to the other ear, representing an event boundary (see dashed yellow line).

Following each sequence, participants completed two temporal memory tests for different pairs of objects: a temporal order test and a temporal distance rating test. Both these tests were designed to assess event separation effects in long-term memory. Because we were specifically interested in changes in subjective temporal memory, we focused on analyzing data from the distance ratings task. The results of the temporal order test are reported elsewhere (Clewett et al., 2025). In the temporal distance ratings task, participants were asked how far apart they remembered two images occurring in the previous sequence, ranging from ‘very close’ to ‘very far’ apart. Importantly, there were always three intervening images (32.5s) between each tested pair, making their objective distance identical. Thus, any differences in distance memory ratings were due to subjective distortions in remembered time.

Replicating prior findings, boundary-spanning pairs were remembered as having been encountered farther apart in time than same-context pairs (*ß* = .071, *SE* = .031, *z* = 2.30, *p* = .022; **Figure 1, bottom right panel**), consistent with the idea that boundaries serve to mentally distance contextually distinct episodes in memory. For more detail on participants’ distance response distributions, see **Supplementary Figure 4**. Interestingly, we also found a significant main effect of pair position (*ß* = .035, *SE* = .0081, *z* = 4.31, *p* < .001), such that later pair positions in the lists were remembered as occurring farther apart in time than item pairs encountered early in the list (see **Supplementary Figure 5**). This pattern suggests a novel time-on-task-like effect for subjective distance ratings as the sequence unfolded.

Additionally, the pitch of the tone changed, as well as the hand required by the participants to make button responses. This pattern continued for the remainder of the 8-item event. After four events and a brief distractor task, participants completed a temporal distance memory test (along with a temporal order memory test discussed elsewhere; Clewett et al., 2025). On each test trial, participants were presented with different pairs of objects from the prior sequence and asked to rate how far apart they thought each pair appeared in time (‘very close’, ‘close’, ‘far’ or ‘very far’), despite their equivalent objective distance. Some tested pairs had been encountered in the same context (in purple), while other pairs had been encountered with an intervening boundary (in yellow). In right panel, raincloud plot displays distance memory ratings for boundary-spanning and same-context pairs. Each dot represents averaged data from one participant, with distance memory ratings (1-4) displayed as a continuous variable. Colored boxplots represent 25^th^-75^th^ percentiles of the data, and the center line represents the median. Statistical significance refers to results from trial-level cumulative link model. \**p* < .05.

### Event boundaries reliably predicted BOLD activation in the VTA, and these brain responses predicted later time dilation effects in memory

In the next analysis, we tested our key hypothesis that event boundaries, or tone switches, engage VTA activation during encoding (see **Figure 2A** for anatomical mask). Consistent with our prediction, we found that Tone Type significantly predicted VTA activation, such that boundary tones elicited higher VTA activation compared to same-context tones (*ß* = 13.45, *SE* = 2.74, *t*(9281.88) = 4.90, *p* < .001). When examining each Tone Type separately, we found that VTA activation was significantly greater than baseline for boundary tones (*ß* = 30.04, *SE* = 7.47, *t*(31.23) = 4.02, *p* < .001), but not for same-context tones (*ß* = 3.35, *SE* = 2.31, *t*(30.29) = 1.45, *p* = .16) (**Figure 2B/C**).

**Figure 2.**
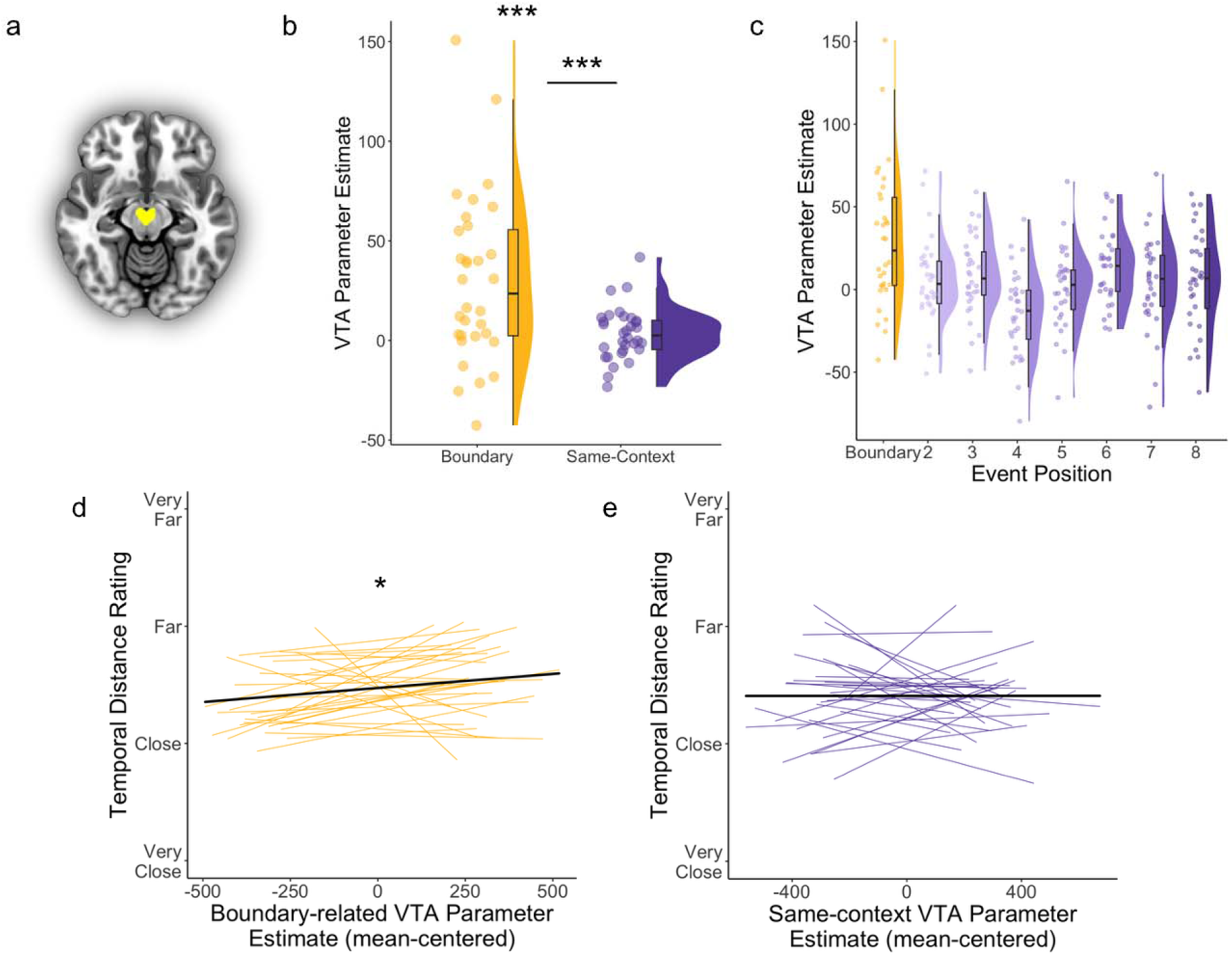
Event boundaries predicted greater VTA activation, and these responses predicted greater time dilation effects in memory. (A) Axial slice of the probabilistic atlas used to identify the VTA (yellow), thresholded at 75% tissue-type probability (Murty et al., 2014). (B) VTA parameter estimates, a measure of brain activation levels, for boundary tones (yellow) and same-context tones (purple). Each dot represents averaged data from one participant. Statistical significance refers to results from trial-level linear mixed effects models. (C) VTA parameter estimates plotted by trial position within the 8-item events during encoding. The first tone in each 8-item event was a boundary trial, when the tone switched from one ear to the other. Same-context tones then played in the same ear and were repeated across item positions 2 through 8 within that stable auditory event. Each dot represents averaged data from one participant. Colored boxplots represent 25^th^-75^th^ percentiles of the data, and the center line represents the median. (D) Participant-level trendlines plotting the relationship between boundary-related VTA parameter estimates (mean-centered within condition) and temporal distance memory ratings (displayed as a continuous variable). Statistical significance refers to results from trial-level cumulative link model. (E) Participant-level trendlines plotting the relationship between same-context VTA parameter estimates and temporal distance memory ratings. Dark, bold lines represent the average linear trend across participants.\*\*\**p* < .001; \**p* < .05.

We next tested if these boundary-related VTA responses predicted increased temporal distance ratings between item pairs spanning those same boundary tones. The results revealed no significant interaction effect of VTA activation and Pair Type (boundary-spanning versus same-context pair) on temporal distance memory (*ß* = .045, *SE* = .032, *z* = 1.40, *p* = .16). However, when analyzing each Pair Type separately, we found that boundary-evoked VTA activation significantly predicted higher temporal distance ratings (*ß* = .10, *SE* = .049, *z* = 2.13, *p* = .034) (**Figure 2D**). In contrast, tone-related VTA activation did not predict temporal distance ratings for the same-context pairs spanning those tones (*ß* = .0030, *SE* = .041, *z* = .073, *p* = .94) (**Figure 2E**). Thus, event boundaries appear to engage the VTA in a way that may distort the temporal structure of memory.

### Event boundaries predicted a momentary increase in blinking

Intriguingly, prior studies have shown that ‘breakpoints’ in experience are associated with transient increases in spontaneous blinking (see Nakano et al., 2009; Wyly et al., 2023), which in turn has been linked to dopaminergic processes (for review, see Jongkees & Colzato, 2016). We sought to replicate these results in the current paradigm by examining how tone switches predict transient changes in blinking (**Figure 3A/B**). This model revealed a main effect of Tone Type on local blink count (*ß* = .12, *SE* = .019, *t*(7461) = 6.07, *p* < .001), such that boundary tones were immediately followed by more blinks than same-context tones (**Figure 3C/D**).

**Figure 3.**
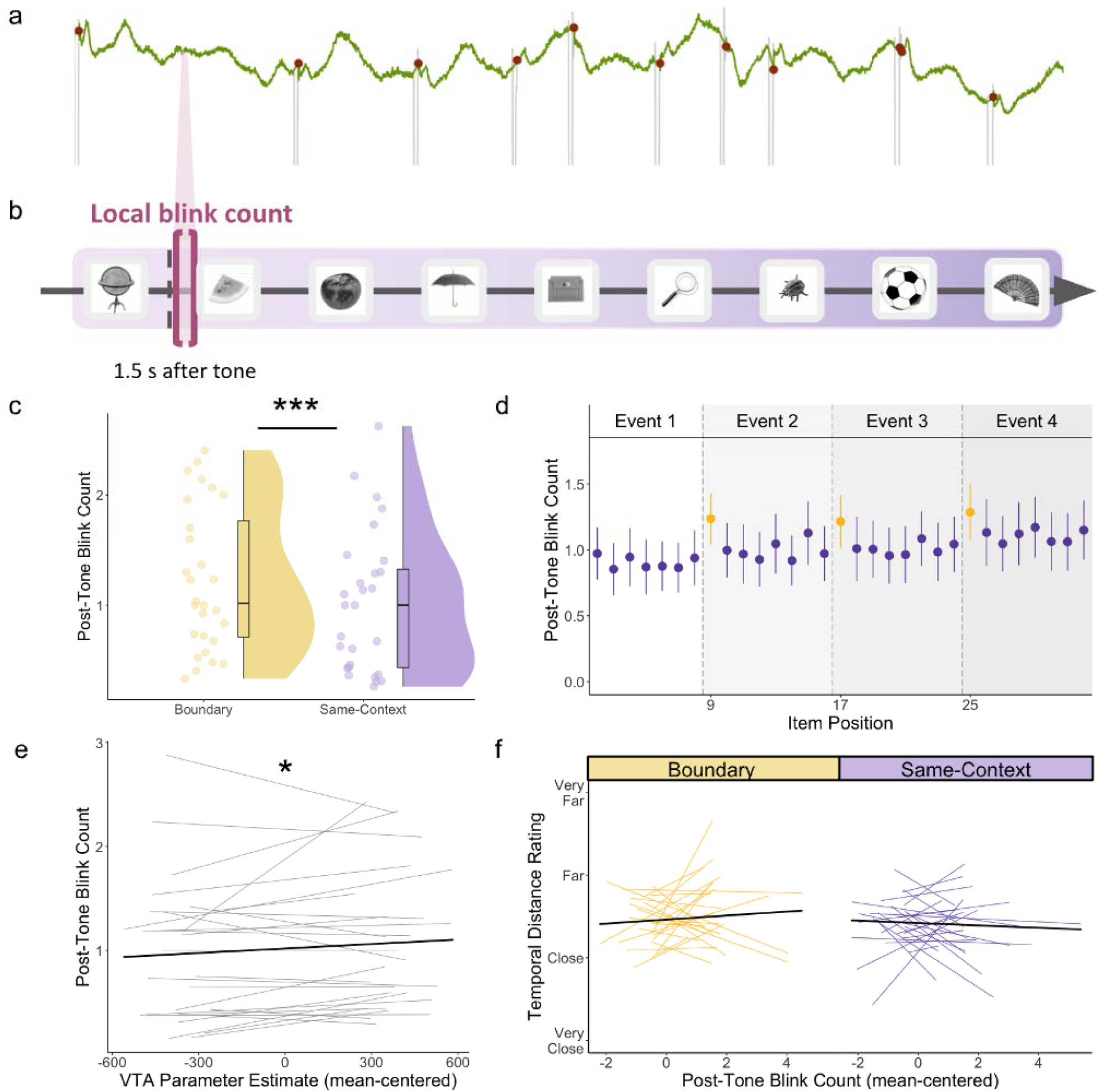
Event boundaries predicted a momentary increase in blinking, and these local changes in blinking were predicted by tone-related VTA activation irrespective of contextual stability or change. (A) Example pupil timeseries for a subset of an encoding block. Green line displays pupil size (in a.u., or arbitrary units) over time. Red circles represent time points when blink artifacts were identified by the pupil preprocessing algorithm based on velocity changes in pupil size. The timeseries is displayed for illustrative purposes and does not strictly align with the trial schemat c. (B) A schematic image of the brief post-tone blink period of interest during event sequence encoding. Local blink count was captured in the short 1.5s interval following the onset of each tone (bracketed in pink). (C) An overall comparison between mean blink count after the onset of boundary tones (yellow) and same-context tones (purple). Each dot represents averaged data from one participant. Statistical significance refers to results from trial-level linear mixed effects model. (D) Mean post-tone blink count by item position in each encoding block (2 through 32; first item excluded). Yellow dots represent blink counts after boundary tones, which always occurred every 8 tones (preceding items 9, 17, and 25). Purple dots represent blink counts after same-context tones, which occurred in all remaining positions in the lists. Error bars represent SEM. (E) Participant-level trendlines plotting the relationship between trial-level VTA parameter estimates (mean-centered) and local post-tone blink count. The dark, bold line represents the average linear trend across participants. Statistical significance refers to results from trial-level linear mixed effects model. (F) Participant-level trendlines plotting the relationship between post-tone blink count (mean-centered) and temporal distance ratings for boundary-spanning pairs (yellow; left) and same-context pairs (purple; right). For modeling, local post-tone blink counts are mean centered for each condition separately. The dark, bold lines represent the average trendline across participants, also for each condition separately. \*\*\**p* < .001; \**p* < .05.

### Tone-related increases in VTA activation predicted blinking irrespective of contextual stability

To validate the putative link between blink behavior and dopaminergic processes, we next examined trial-level coupling between post-tone blink count and tone-related VTA activation. We found a significant main effect of VTA activation on post-tone blink count (*ß* = .00025, *SE* = .00012, *t*(7270) = 2.08, *p* = .038), such that greater tone-related VTA activation was associated with higher blink count irrespective of condition (**Figure 3E**). There was no significant VTA-by-tone type interaction effect on post-tone blink count (*ß* = .00016, *SE* = .00012, *t*(7271) = 1.34, *p* = .18), reinforcing a more general, context-invariant association between phasic VTA activation and blinking.

To verify that blink changes likely resulted from tone-related VTA activation (and not vice versa), we also performed a control analysis focusing on blinks that occurred within the 1.5s period *before* each tone. Pre-tone blink count was not significantly predicted by Tone Type (*ß* = .017, *SE* = .020, *t*(7410) = .83, *p* = .41) or tone-related VTA activation (*ß* = .00012, *SE* = .00013, *t*(7224) = .95, *p* = .34). Together, these blink timing-related control analyses provided strong evidence that blinking was triggered by behaviorally relevant changes during a dynamic experience.

### Momentary increases in local boundary-evoked blinking behavior were not associated with later temporal distance ratings in memory

Finally, we tested if post-tone blink count was related to later memory separation effects. Contrary to our prediction, there was no main effect of post-tone blink count (*ß* = .019, *SE* = .038, *z* = .50, *p* = .62) or blink-by-pair type interaction effect on temporal distance ratings (*ß* = .060, *SE* = .039, *z* = 1.54, *p* = .12; **Figure 3F**).

### Testing relations between more prolonged periods of blinking, VTA activation, and temporal distance memory

Beyond local changes in blinking around boundaries, we were also interested in testing whether blinking across larger timescales (i.e., windows spanning approximately 32.5 seconds between to-be-tested item pairs) predicted distance memory and VTA activation (**Figure 4A/B**).

**Figure 4.**
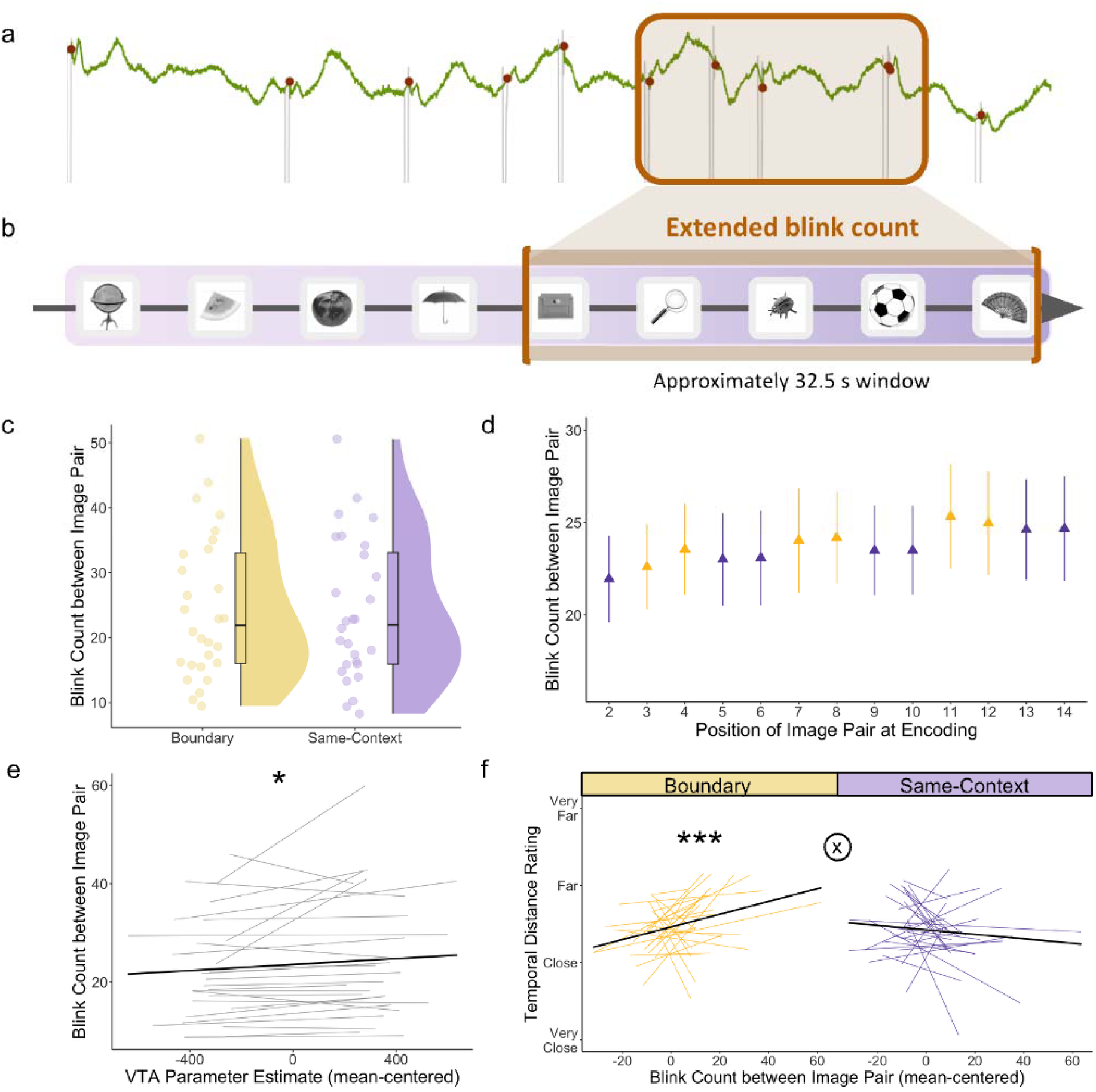
Event boundary-evoked VTA activation predicted greater sustained blinking between to-be-tested item pairs, and this blink pattern predicted later time dilation specifically between boundary-spanning item pairs in memory. (A) Example pupil timeseries for a subset of an encoding block. (B) Schematic of blink window of interest during event sequence encoding. Extended blink count was captured during each 32.5s window between the onset and offset of to-be-tested item pairs (bracketed in orange). (C) An overall comparison between mean blink count between to-be-tested images spanning an event boundary (yellow) or from the same context (purple). Each dot represents averaged data from one participant. (D) Mean blink count during these longer time windows, plotted by item pair position at encoding (14 tested pairs total; first pair excluded). Yellow triangles represent boundary-spanning item pairs, while purple triangles represent same-context item pairs. In order of pair presentation at encoding, the boundary-spanning pairs were the 3rd, 4th, 7th, 8th, 11th, and 12th item pairs. Error bars represent SEM. (E) Participant-level trendlines plotting the relationship between trial-level VTA parameter estimates (mean-centered) and more sustained blinking patterns between to-be-tested item pairs. The dark, bold line represents the average linear trend across participants. Statistical significance refers to results from trial-level linear mixed effects model. (F) Participant-level trendlines plotting the relationship between extended blink count (mean-centered) and temporal distance ratings for boundary-spanning pairs (yellow; left) and same-context pairs (purple; right). For modeling, blink count was mean-centered for each condition separately. The dark, bold line represents the average trendline across participants, also for each condition separately. Statistical significance refers to results from trial-level cumulative link models. The circled “X” symbol represents a significant condition-by-blink count interaction effect on temporal distance ratings with *p* < .01. \*\*\**p* < .001; \**p* < .05.

### Increased blinking between to-be-tested item pairs was related to tone-related VTA activation and time dilation effects in memory

First, we tested whether boundaries modulated blink count across these larger time periods. We found that Pair Type (boundary-spanning pair versus same-context pair) did not significantly predict blink count across the to-be-tested pair windows (*ß* = .26, *SE* = .17, *t*(2881.02) =1.55, *p* = .12; **Figure 4C/D**).

However, tone-related VTA activation significantly predicted blink count in the surrounding time window (*ß* = .0030, *SE* = .0010, *t*(2795) = 2.94, *p* = .0033), such that greater tone-related VTA activation predicted higher blink count in general (**Figure 4E**). Like more local changes in blink count, there was no significant VTA-by-pair type interaction effect on sustained blinking behavior (*ß* = .00048, *SE* = .0010, *t*(2796) = .46, *p* = .64), suggesting that momentary VTA activation is coupled with prolonged blinking patterns more generally. Thus, despite differences in their timescales of action, momentary increases in VTA activation were meaningfully embedded within sustained periods of elevated blinking across sequence encoding.

We also found a significant blink-by-pair type interaction effect on temporal distance ratings (*ß* = .012, *SE* = .0039, *z* = 3.17, *p* = .0016). When analyzing the two pair types separately, we blink count significantly predicted temporal distance ratings for boundary-spanning item pairs (*ß* = .018, *SE* = .0058, *z* = 3.16, *p* = .0016), but not for same-context item pairs (*ß* = -.0072, *SE* = .0053, *z* = −1.38, *p* = .17; **Figure 4F**). This finding suggests that meaningful context shifts predicted stronger coupling between prolonged blinking patterns and the amount of subjective time encoded into long-term memory.

### Identifying which blink periods and stimuli predicted subsequent temporal distortions in memory

The results so far demonstrate that transient blink count in the 1.5 sec post-tone-switch did not predict temporal distance ratings between object pairs. Yet, the longer period of blinking between to-be-judged items did predict temporal distance ratings across an event boundary. Because the momentary boundary-related increase in blinking was encompassed within the larger temporal interval between pairs, it is likely the case that blinking in some other time part of the interval (i.e., not specifically within the 1.5s after the tone switch) was contributing to the observed relationship to time dilation. To explore this nuance of the data, we investigated which specific periods of extended blinking drove the time dilation effect. In brief, we found that the aggregation of tone-evoked blinking during the later portion of the inter-pair interval (i.e., from the tone switch to the appearance of the second to-be-judged item) predicted significant time dilation effects across boundaries (*ß* = .036, *SE* = .017, *z* = 2.11, *p* = .035). For more details about these blink analyses and results, see the **Supplementary Material** and **Supplementary Figure 12**.

### Testing the specificity of blink and distance memory measures to the dopaminergic system

Recent findings from this same dataset showed that pupil-linked arousal and locus coeruleus (LC), a key hub of the brain’s arousal and noradrenergic system, were associated with impairments in temporal memory (Clewett et al., 2025). An important open question is whether our current findings on subjective temporal memory and blinking are linked to arousal-related neuromodulation more generally. According to theoretical frameworks of event segmentation (Zacks & Sargent, 2010), both the dopaminergic and noradrenergic systems are well positioned to elicit a global updating signal when event transitions occur. However, it is possible that they contribute to memory separation (Rouhani et al., 2024) and blink behavior in different ways. We have also previously shown that LC activation at boundaries relates to more differentiated multivoxel activation patterns in left dentate gyrus (DG), a hippocampal subregion that is important for memory encoding and in disambiguating overlapping neural representations (Clewett et al., 2025). Much work shows that dopaminergic modulation of hippocampal processing also plays an essential role in structuring and encoding episodic memories, making it another candidate input signal for memory separation at its target regions (Shohamy & Adcock, 2010).

To test the specificity of dopaminergic effects on temporal and hippocampal processing, we ran several mixed effects modeling analyses that complemented those in Clewett et al. (2025), which had focused only on LC-related effects. We then also performed follow-up modeling to test whether relationships between VTA activation, eye-tracking measures (pupil dilation and blinking), hippocampal pattern stability, and temporal memory differed from those of the LC (see **Supplementary Material** for details on methods and results). In brief, we found that LC activation was more strongly coupled with impairments in temporal order memory, an objective behavioral index of memory separation, than VTA activation. However, VTA activation was not significantly more coupled with distance ratings than LC activation, so we cannot make strong claims about specificity. On the other hand, blinking was more strongly coupled with VTA activation than LC activation, and more strongly coupled with distance memory than impairments in order memory.

When probing links to the stability of hippocampal multivoxel patterns across time, LC activation was significantly more coupled with pattern differentiation in left DG compared to VTA activation. This suggests that noradrenergic processes may have a stronger influence over ongoing event representations supported by the hippocampus. Finally, VTA activation was not significantly correlated with different features of boundary-evoked pupil dilation (see Clewett et al., 2025). For more details concerning all these analyses and results, see **Supplementary Material and Supplementary Figures 6-11.**

Taken together, our results suggest that extended blinking behavior is indeed closely tied to dopaminergic activity as opposed to arousal processes more generally. However, they suggest that the noradrenergic system, and not the dopaminergic system, may modulate memory separation by influencing objective aspects of temporal memory and differentiating hippocampal representations.

## Discussion

Segmenting continuous experience into distinct events helps structure memories in ways that support adaptive behavior. One way this structure might be achieved is through the subjective ‘stretching’ of remembered time across an abrupt change in context, such as a shift in location, goal, or mood (for review, see DuBrow, 2024; Clewett et al., 2019, 2020; Ezzyat & Davachi, 2014; Cowan et al., 2024). Although these subjective distortions can belie the actual passage of time, they may serve a functional role in delineating boundaries between events in both perception and memory. In the current neuroimaging study, we replicated this time dilation effect in memory and found that context shifts, or event boundaries, elicited transient increases in ventral tegmental area (VTA) activation – a key hub of the dopamine system – and blinking. VTA activation was associated with both momentary and sustained increases in blink count, supporting a putative link between blink behavior and dopaminergic activity. Finally, stronger VTA responses and elevated blink counts across boundaries predicted greater time dilation between item pairs spanning those same boundaries. Together, these converging results suggest that dopaminergic signaling may contribute to the mnemonic separation of distinct events by expanding the temporal gap between them.

Our findings help bridge separate lines of research linking dopamine to episodic memory and temporal processing. We first showed that context shifts predicted stronger activation of the VTA, consistent with prior work showing that dopamine flags novel and salient stimuli (Duszkiewicz et al., 2019; Shohamy & Adcock, 2010). Context shifts also predicted brief increases in blinking, aligning with studies showing that blinks tend to occur near event transitions, including punctuation marks in text passages or breakpoints in everyday conversations (Hall, 1945; Wyly et al., 2023; Nakano et al., 2009; but also see Nakano et al., 2013 for an alternative interpretation). Specifically, we found that these blinks occurred after boundary tones, not before them. This timing is consistent with evidence that blinking tends to be delayed when a motor response is required (Demiral et al., 2023), such as a button press response to an orienting question (“Is this object larger or smaller than a standard shoebox?”). Moreover, VTA responses to tones generally predicted transient increases in blinking. This finding aligns with recent evidence that discrete blinks are associated with fMRI activation in dopaminergic regions, including the VTA and substantia nigra (Demiral et al., 2023). Overall, these results support the idea that VTA activation and blinks mark transition points in experience.

Why might dopaminergic processes be engaged at event boundaries? One possibility is that dopamine facilitates the rapid updating of event models during sudden or unexpected changes in the environment. Supporting this view, prior neuroimaging work shows that dopaminergic brain regions – such as the substantia nigra and basal ganglia – are activated when participants make predictions about the next few seconds of videos depicting everyday activities (e.g., gardening). When these intervals contain an event boundary, participants tend to make less accurate predictions (Zacks et al., 2011). Dopamine may signal high prediction error at event boundaries, indicating that a new mental representation of the world is needed to represent unfolding events more accurately (Zacks et al., 2011; also see Zacks et al., 2007). It has also been shown that VTA activation relates to moments of surprise in more naturalistic settings, such as watching videos of basketball games (Antony et al., 2021). However, given that the structure of our task was largely predictable, we speculate that it is unlikely that prediction errors and surprise accounted for boundary-related patterns of VTA activation in the current sequence paradigm.

An alternative, more plausible explanation for boundary-evoked dopaminergic activity is that it signals contextual novelty during dynamic experience. It has been theorized that the VTA releases dopamine into the hippocampus in response to ‘common novelty’ – that is, stimuli that are novel under current conditions but not completely new (Duszkiewicz et al., 2019). Following this logic in the current study, each context change was relatively novel within its local temporal context (i.e., block), because the tone switches occurred infrequently and introduced a pitch that was unique to a given block. Thus, dopamine might signal a kind of common novelty at these transitions within sequences, marking the end of one meaningful event and the start of the next.

Consistent with our key hypothesis, we also found that higher boundary-related VTA activation predicted greater time dilation effects in subsequent memory, with individuals remembering item pairs as being encountered farther apart in time if they spanned a boundary. This finding aligns with prior research showing that dopaminergic processing relates to changes in internal representations of time. For example, studies in humans have found that encountering salient oddball stimuli, linked to dopamine signaling (Bunzeck & Düzel, 2006), can lead individuals to overestimate the duration of those stimuli (see Failing & Theeuwes, 2016; Gupta, 2022; Ongchoco, Wong, & Scholl, 2023). However, we note that this effect of VTA activation on time dilation did not differ between conditions, so we cannot draw strong conclusions about specificity. It is possible that this VTA-memory relationship reflects processes beyond boundary-specific signaling, such as general salience or updating mechanisms.

Nevertheless, because the coupling between boundary-evoked VTA activation and temporal memory emerged at the trial level, our results suggest that transient dopaminergic responses at event boundaries might meaningfully shape how temporal distance is encoded. We also note that interpreting VTA-memory relationships for same-context pairs is inherently constrained by choosing a single tone to index VTA activation: the reference tone that was position-matched to the tone switch between boundary-spanning pairs. It remains possible that there are activity patterns at other timepoints within this same-context pair interval that could yield a different pattern of results.

The time dilation effect we observed aligns with existing conceptual frameworks of temporal processing, including the dopamine clock hypothesis and contextual-change hypothesis of remembered duration. According to these models, both dopamine (Meck, 1983; 1996; see Fung, Sutlief, & Hussain Shuler, 2022) and the number of contextual changes stored in memory (Block, 1992, Block & Zakay, 1996) serve to expand internal representations of time. One intriguing possibility is that boundary-induced VTA activation and dopamine release boost encoding of local context changes, constructing more memories of salient experiences. As a result, there is a greater amount of episodic information and details available at retrieval, leading to overestimations of retrospective time. Future work could test this idea by examining whether boundary-induced VTA activation relates to increases in local source encoding processes, as boundaries have been shown to reliably enhance source memory for concurrent information (Clewett et al., 2020; Cowan et al., 2024; McClay et al., 2023). We suggest that these dopamine-mediated temporal distortions may occur via modulation of hippocampal activity, given its well-known role in supporting the storage and retrieval of temporal and source information (see Howard & Eichenbaum, 2013).

On a larger timescale, we observed that prolonged periods of blinking across boundaries predicted remembered time. This relationship was stronger for periods spanning contextual change versus contextual stability, suggesting that boundary-specific features like contextual novelty and task relevance might be critical for shaping memories of time in a meaningful way. Indeed, our analytical approach to the blink data was novel in that prior eye-tracking studies have either measured discrete blinks after individual stimuli (e.g., Terhune, Sullivan, & Simola, 2016) or longer periods of spontaneous blinking offline or during rest. For example, spontaneous blink rate has been linked to dopaminergic processes in non-human primates and humans with neuropsychiatric disorders (for review, see Jongkees & Colzato, 2016). Resting-state measures of spontaneous blinking are likely noisy, given that blinking is affected by both cognitive and non-cognitive factors, such as fatigue. The current study assessed blinking during a behaviorally relevant time window during memory encoding, perhaps leading to a stronger and more meaningful coupling between blinking and dopaminergic activity. One possibility, then, is that blinking may better track dopamine dynamics online and over longer periods of time, potentially reconciling some of the mixed findings in the blink-dopamine literature (see Blin et al., 1990; Demiral et al., 2022, 2023; Jongkees & Colzato, 2016; Strakowski et al., 1996; Strakowski & Sax, 1998).

There are several limitations in the current study that warrant consideration. First, fMRI cannot establish a causal link between dopaminergic processing and time dilation in memory, nor can it directly assess neurotransmitter activity. Second, while we used a highly specific, data-driven algorithm to locate blink artifacts, examining electrical activity near the eye offers a more direct method of detecting blinks (Cruz et al., 2011). Third, there are mixed findings concerning the existence or strength of an association between blinking and dopaminergic processing. Some studies challenge the relationship between certain measures, such as striatal dopamine synthesis capacity, and blinking (Sescousse et al., 2018; van den Bosch et al., 2023; also see Dang et al., 2017). Additional factors that could influence blink behavior include the activity of other neuromodulators like serotonin (see e.g., Gunduz et al., 2024), as well as inattentiveness or fatigue (see Maffei & Angrilli, 2018). However, because event boundaries typically elicit an increase in attention (Heusser et al., 2018), it is unlikely that the effects we observed were solely or predominantly driven by fatigue or time-on-task effects (Stern et al., 1994; Maffei and Angrilli, 2018). Future studies should examine how relations between blinking and cognitive processing may be altered under different conditions, including whether blinking is measured online or during rest.

In summary, our findings revealed that different indirect markers of dopaminergic processing are related to time dilation effects in memory. We speculate that these subjective time distortions are highly adaptive, because they might serve to increase the representational distance between adjacent episodes in memory. These results are an important first step towards unraveling the complex role of dopamine activity in shaping everyday memory organization. Still, additional techniques are needed to assay neuromodulatory activity more directly and at a higher temporal resolution (e.g., Bang et al., 2023). Ultimately, these findings will promote a deeper understanding of how internal representations of time mold the complexity of our experience into a meaningful record of past events.

## Methods

### Participants

Thirty-six participants were recruited from the New York University (NYU) Psychology Subject Pool and the local community. Sample size was estimated using a G*Power 3.1 power analysis (alpha = 0.05, power = 0.80, *d* = 0.80), based on the pooled temporal memory performance from a very similar event boundary experiment (Clewett, Gasser, & Davachi, 2020). This power analysis yielded a required sample size of 29 participants. To account for attrition, we recruited 36 participants.

Participants were required to have normal or corrected-to-normal vision and hearing, to have no metal in their body, and to not be taking beta-blockers or psychoactive medications. All participants provided written informed consent approved by the NYU Institutional Review Board and received monetary compensation for participation.

Four participants were fully excluded from analyses (reasons included falling asleep in the scanner and audio malfunction). Therefore, our final sample size was 32 participants (20 F; *M*_age_ = 22 years, *SD*_age_ = 2.7 years). Of this sample, 15 participants identified as Asian, 2 as Black/African American, 4 as more than one race, and 11 as White.

Of these 32 participants, five requested to leave the scanner early and thus did not complete all 10 blocks of the task (*n* = 2 completed 9 blocks, *n* = 1 completed 8 blocks, *n* = 2 completed 7 blocks). Finally, four participants’ eye tracking data was excluded due to equipment malfunction or poor quality, leaving 28 participants with usable data for all analyses that included blink data.

### Materials

Participants viewed 512 grayscale images of everyday objects (Gabrieli et al., 1997; Kensinger et al., 2006), which were resized to 300 x 300 pixels, placed on a gray background, and luminance-normalized using the MATLAB SHINE toolbox to control for the effect of brightness on pupil behavior. To embed these images within distinct auditory contexts, participants were presented with 1s pure tones (created with Audacity; https://www.audacityteam.org/) in either the left or right ear. The six tone frequencies used ranged from 500-1000 Hz in 100 Hz intervals, were perceptually discriminable, and were sufficiently arousing to facilitate engagement in the task.

### Code and data accessibility

The code, experiment materials, and data for this study will be made publicly available on the OSF account of E.M. (osf.io/yt6hm) upon acceptance of this manuscript.

### Overview of experimental design

This experiment took place over two days. On the first day, participants completed several questionnaires and a practice task block before completing the full event sequence task in the MRI scanner. The MRI session took approximately 3 hours, with 2.5 hours of scanning. On the second day, approximately 24 hours later, participants returned to perform a surprise item recognition memory test (not described in the current paper).

### Event sequence encoding task

While in the MRI scanner, participants completed a modified version of an event sequence encoding task that has been shown to elicit reliable event segmentation effects in memory, as indexed by significant time dilation in retrospective distance ratings (Clewett et al., 2020) (**Figure 1**). Our goal was to use these boundary-induced subjective temporal distortions to operationalize whether memories had become separated at context shifts. That is, if item pairs spanning a boundary were remembered as appearing farther apart than item pairs from the same context – despite being the same objective distance apart during encoding – then they were more likely to have become encoded as distinct episodic memories.

In this encoding-retrieval block design, participants were presented with sequences of 32 object images for 2.5s each. For each image, participants made a button press to respond to a simple orienting question (“Is this object larger or smaller than a standard shoebox?”). Additionally, participants were asked to form their own mental narrative of the image sequence to encourage associative encoding (e.g., DuBrow and Davachi, 2013). A jittered ISI (3, 5, or 7s fixation cross) separated each image presentation to optimize deconvolution of the blood-oxygen level dependent (BOLD) signal during the fMRI analyses.

Each item sequence was divided into four auditory ‘events’ that contained 8 images each. Events were defined by a stable auditory context, in which 1s pure tones played either in participant’s left or right ear only, halfway through each jittered ISI. The laterality of the tone also cued participants to use either their left or right hand to make button responses for the object size judgments (left ear = left hand).

Thus, the tones not only provided sensory information to elicit perception of stable mental events, but were also task relevant. After the final image in an 8-item event, the tone abruptly switched sides to the opposite ear and changed pitch. Participants were also instructed to switch the hand used to make button responses. This salient change constituted an ‘event boundary’ in the sequence, marking the end of one event and the beginning of the next. The new context then remained the same for the rest of the 8-item event.

Each item sequence contained three tone switches total, resulting in four stable auditory events. There were 10 encoding-retrieval blocks total across the experiment. The starting ear/hand was counterbalanced across blocks. Pitch changes were pseudorandomized, such that the same frequency (e.g., 500 Hz) was not heard more than once in any given encoding block. To separate the encoding and retrieval tasks and to reduce any recency effects in memory, participants completed a brief arrow detection task (45s) after each encoding block. Participants were presented with 0.5s left-facing (<) or right-facing (>) arrows in the center of the screen, separated by 0.5s ISIs of fixation. They were instructed to indicate which direction the arrow was facing via button press as quickly as possible.

### Temporal distance memory test

At the end of each encoding block, participants completed two temporal memory tests: the first for temporal order and the second for temporal distance. Given that our primary focus was on subjective distortions in distance memory, the results from the temporal order memory test are reported elsewhere (Clewett et al., 2025).

In the temporal distance memory test, 14 item pairs from the encoding sequence were presented for a fixed duration of 5s each. Importantly, each of these item pairs had been separated by exactly three images during encoding. Because the objective distance between to-be-tested pairs was pseudorandomized to always be equivalent (32.5s from onset of the first image to the offset of its pairmate), we were able to measure subjective temporal distortions in memory across conditions.

Accordingly, participants were asked to judge how far apart in time two images appeared. There were four options: ‘very close,’ ‘close,’ ‘far,’ and ‘very far.’ To replicate previous effects of event boundaries on temporal distance memory, we compared ratings for two types of item pairs: same-context pairs (8 trials per block) and boundary-spanning pairs (6 trials per block). Same-context pairs contained images from the same context during encoding, while boundary-spanning pairs contained images that spanned a change in context (i.e., task-relevant tone switch) during encoding.

We excluded the first same-context item pair from all analyses, because this pair contained the first image in each block. As such, it likely constituted a task-irrelevant event boundary and would therefore produce different behavioral effects than the other same-context pairs. Due to a programming error, one of the boundary pairs from each list also contained an incorrect item. Data for these specific pairs (appearing once per block across 23 participants) were excluded from the analyses. Finally, one block was excluded for 9 participants due to a timing error.

### fMRI Acquisition and preprocessing

#### fMRI/MRI data acquisition

All neuroimaging data were acquired with 3T Siemens Magnetom PRISMA scanner using a 64-channel matrix head coil. First, participants underwent a high-resolution MPRAGE T1-weighted anatomical scan (slices = 240 sagittal; TR = 2300ms; TE = 2.32ms; TI = 900ms; FOV = 230 mm; voxel in-plane resolution = 0.9 mm^2^; slice thickness = 0.9 mm; flip angle = 6°; bandwidth = 200 Hz/Px; GRAPPA with acceleration factor = 2; scan duration: 5m 21s).

Next, participants underwent a T2-weighted functional scan (slices = 240 sagittal; TR = 3200ms; TE = 564 ms; FOV = 230 mm; voxel in-plane resolution = 0.9 mm^2^; slice thickness = 0.9 mm; flip angle = 6°; bandwidth = 200 Hz/Px; GRAPPA with acceleration factor = 2; scan duration: 3m 7s). The task audio was also calibrated during this scan to ensure the tones were audible above scanner noise and could be discriminated from one another. Participants could make button presses to request changes in tone volume. Additionally, we collected two fieldmap scans to assist with functional imaging unwarping (1 in anterior-posterior (AP) phase encoding direction and 1 in posterior-anterior (PA) phase encoding direction). Afterwards, we collected a short fast spin echo (FSE) sequence MRI scan that enables visualization of the locus coeruleus (LC). Details are reported elsewhere (Clewett et al., 2025).

After the structural scans were collected, participants underwent separate functional imaging for each of the 10 encoding blocks and the 10 retrieval blocks. Functional scans were collected using a whole-brain T2*-weighted multiband echo planar imaging (EPI) sequence (128 volumes per encoding block; TR = 2000ms; TE = 28.6ms, voxel in-plane resolution = 1.5 x 1.5 mm^2^; slice thickness = 1 mm with no gap; flip angle = 75°, FOV = 204 mm X 204 mm; 136 X 136 matrix; phase encoding direction: anterior-posterior; GRAPPA factor = 2; multiband acceleration factor = 2). In each volume, 58 slices were tilted - 20° of the anterior commissure-posterior commissure line and were collected in an interleaved order. A single-band reference image was collected for each run of the task, but was not included during preprocessing.

#### fMRI preprocessing

Image preprocessing was performed using FSL Version 6.00 (FMRIB’s Software Library, www.fmrib.ox.ac.uk/fsl). Functional images were preprocessed using the following steps: removal of non-brain tissue using BET; B0 unwarping using fieldmap images; grand-mean intensity normalization of the 4D data set by a single multiplicative factor; and application of a high-pass temporal filter of 100s. No spatial smoothing was applied due to the small size of the VTA and to preserve spatial specificity. Motion correction was performed using the MCFLIRT tool, which produced six motion nuisance regressors. Additionally, fsl_motion_outliers was used to identify volumes with extreme head movements, or frame displacements, using the DVARS option. Both this matrix of outlier volumes and the six motion regressors were as modeled as covariates in the subsequent GLM analyses. Entire blocks with excessive head motion overall (conservatively defined as mean frame displacement > 1 mm) were excluded from analysis, resulting in the removal of one block each from three participants. Each participant’s denoised mean functional volume was co-registered to their T1-weighted high-resolution anatomical image using brain-based registration (BBR). Anatomical images were then co-registered to the 2 mm isotropic MNI-152 standard-space brain using an affine registration with 12 degrees of freedom.

#### Physiological denoising

Eight separate physiological nuisance signal regressors were extracted for the subsequent GLM analyses. First, FSL FAST was used to decompose each participant’s high-resolution anatomical images into probabilistic tissue masks for white matter (WM), grey matter (GM), and cerebrospinal fluid (CSF). The CSF and WM masks were thresholded at 75% tissue-type probability to increase their spatial specificity and reduce potential overlap. Following a similar approach to Bartoň et al. (2019), we defined eight 4 mm spheres in representative regions of WM and CSF (four of each type; for exact coordinates, see Bartoň et al., 2019). The eight spheres and WM and CSF anatomical masks were then transformed into each participant or block’s native functional space and merged to further increase their spatial specificity. Nuisance timeseries for each of the four WM and four CSF merged masks were then extracted from each block’s preprocessed functional data and modeled as nuisance regressors in the GLMs.

#### VTA region-of-interest (ROI) definition

To create an anatomical mask of the VTA, we used a publicly available probabilistic atlas that was originally created using individual hand-drawn ROIs (Murty et al., 2014). This standard-space VTA mask was thresholded at 75% probability to increase its spatial specificity and then registered into each participant’s functional run (i.e., block) of the encoding task.

### fMRI analyses

#### Generalized linear modeling (GLM) analyses and acquisition of single-trial activation estimates of VTA activation

To estimate VTA activation at the trial level, we conducted Least Squares Separate (LSS) GLM analyses on the unsmoothed functional data from the sequence encoding task. This modeling approach generates unique activation estimates for each stimulus (i.e., tone or image) from the task. Importantly, the repetition or switching of tones carried the critical information about the stability or change in the surrounding context. Thus, using LSS GLM enabled us to isolate the distinct effects of context-relevant tones on brain activation.

Each LSS-GLM contained a total of 64 task-related regressors for each block of the encoding task, because there were 32 tones and 32 images in each block. Each tone was modeled as a 1s stick function and each image was modeled as 2.5s stick function. Both regressors were convolved with a dual-gamma hemodynamic response function (HRF). A total of 64 separate LSS-GLM analyses were conducted for each block of the ten blocks of the task, where one stimulus (image or tone) served as the regressor of interest and all other trials were modeled as a separate regressor. Each LSS-GLM thereby resulted in a unique activation estimate (i.e., beta map) across the whole brain for each stimulus (Mumford et al., 2012; Mumford, Davis, & Poldrack, 2014). To control for noise, a total of 14 nuisance regressors (4 WM, 4 CSF, and 6 motion regressors) were included in each GLM, along with individual nuisance regressors for trials with extreme head movements.

A VTA ROI analysis was performed on the resulting beta maps. The standard-space VTA anatomical mask was transformed into each participant’s native run-space for each encoding block and used to extract single-trial VTA parameter estimates (a measure of activation) for each of the 32 tone trials (3 boundary tones, or switches, and 29 same-context tones, or repeats). We excluded noisy datapoints using boxplot outlier removal by participant, resulting in the removal of approximately 3% of the tone-related VTA activation trials across the entire dataset.

### Eye-tracking methods

#### Eye-tracking

Pupil diameter was measured continuously at 250 Hz during the event sequence task using an infrared EyeLink 1000 eye-tracker system (SR Research, Ontario, Canada). Raw pupil data, segmented by block, were preprocessed using ET-remove-artifacts, a publicly available MATLAB program (https://github.com/EmotionCognitionLab/ET-remove-artifacts). Although this software is typically used to clean pupil timeseries by interpolating over blink events and other artifacts (e.g., Clewett et al., 2025), our primary objective was to identify blink event timestamps and quantify their frequency within different periods of interest.

ET-remove-artifacts located blink events by identifying rapid changes in pupil size, or pupil velocity, following the approach described in Mathôt et al. (2013). The velocity timeseries was computed by applying MATLAB’s finite impulse response (FIR) differentiator filter on the raw pupil size timecourse. This method provides a robust estimate of instantaneous rate of change while minimizing noise amplification. For our dataset, we selected FIR filter parameters (Filter Order = 4, Passband Frequency = 10, and Stopband Frequency = 12) to produce a smooth velocity timeseries with trough-and-peak profiles that were identifiable and specific to blink events rather than noise-related data loss.

The algorithm then used MATLAB’s “findpeaks” function to locate peaks and troughs in the pupil velocity timecourse. Blink profiles were distinctly identifiable as a contiguous trough followed by a peak in the velocity timeseries. To achieve a high degree of specificity to blink events, we set the Peak and Trough Threshold Factor and Trough Threshold Factor at 8 standard deviations of the velocity timeseries. This threshold is higher than those typically used for cleaning artifacts from pupil timeseries (e.g., Clewett et al., 2025), ensuring that only substantial blink-related troughs and peaks were counted as blink events rather than small and transient velocity changes attributable to other artifacts.

To generate a preprocessed pupil timecourse, the algorithm applied a linear interpolation across identified blink intervals. Artifact intervals exceeding 2 seconds were automatically imputed with NaN (missing data indicator).

Alongside the blink estimates, we also computed pupil dilation responses to both boundary tones and same-context tones. A temporal principal component analysis (PCA) was applied to these tone-evoked pupil responses to dissociate distinct autonomic and functional components of pupil-related arousal. Details about pupil analyses and methods are reported elsewhere (Clewett et al., 2025), because the goal of this manuscript was to focus on the specificity of the relationship between brainstem nuclei activation and eye-tracking measures of neuromodulation. As mentioned previously, four participants were excluded from eye tracking analyses due to system malfunction or poor overall eye tracking quality, resulting in a total of 28 participants for all eye tracking-related analyses.

#### Computing local blink counts

Breakpoints in continuous experience have been shown to elicit a momentary increase in blinking (see Nakano et al., 2009; Wyly et al., 2023). Further, blink behavior has also been putatively linked to dopaminergic processes (Jongkees & Colzato, 2016), suggesting that blink behavior may offer a window into neuromodulatory processes that facilitate memory separation at boundaries. To test this idea, we tabulated local blink count in a 1.5s interval immediately after the onset of each tone. The size of this interval was chosen because the smallest ISI in the current paradigm was 3s – that is, 1.5s on either side of the tone onset. In this way, we were able to capture tone-induced blinks during a brief period that was ‘uncontaminated’ by the preceding or ensuing images on any trial. To acquire highly conservative estimates of blink behavior that were not confounded by noise or data loss, blink intervals with more than 25% invalid samples were excluded from analyses. As with the VTA data, we also cleaned the trial-level blink responses using boxplot outlier detection within each participant. After these exclusions, approximately 78% of local blink intervals remained per participant, on average.

#### Computing sustained blink counts between to-be-tested item pairs

We also zoomed out to examine blink patterns across longer, behaviorally relevant windows of encoding to assess more sustained dopaminergic processes. Blink count was calculated for the 32.5s window between each to-be-tested pair, from the onset of the first item to the offset of its pairmate (encountered four images later). We conducted the same noise-related and by-participant boxplot outlier removal procedure as before, which left approximately 79% of blink windows per participant, on average.

### Testing linear relations between the VTA, temporal memory, and blink patterns

To assess trial-level relationships between our key variables, we used linear mixed-effects and cumulative link modeling. Linear mixed-effects modeling was used when the outcome of interest was continuous (i.e., trial-level VTA parameter estimates, post-tone blink count, and extended blink count). Fixed effect predictors in these models were categorical (i.e., tone type: boundary tone, same-context tone) and/or continuous (i.e., trial-level VTA parameter estimates).

Cumulative link models are the most common type of ordinal regression model, using a similar frequentist approach to linear mixed-effects models (Christensen, 2015; Dunn, 2020). Thus, cumulative link modeling was used when the outcome of interest was ordinal (i.e., trial-level temporal distance memory ratings). Distance memory ratings were ordered from the smallest to largest rating: ‘very close,’ ‘close,’ ‘far,’ ‘very far.’ Fixed effect predictors in these models were categorical (i.e., pair type: boundary-spanning pair, same-context pair) and/or continuous (i.e., trial-level parameter estimates, post-tone blink count, and extended blink count). In the models with trial-level VTA parameter estimates as a fixed-effect predictor of distance memory, we included an additional categorical variable for Pair Position during encoding to account for potential list position effects. Including this predictor improved model fit (*p*s < .01). This predictor was also included in associated modeling analyses in the **Supplementary Material** (see **Supplementary Figures 6-7**).

To meaningfully relate local measures (i.e., VTA parameter estimates and post-tone blink counts) to measures capturing a wider temporal window (i.e., extended blink count and distance memory ratings), we extracted the value associated with one tone between each to-be-tested item pair. For details on the exact positions of the selected tones and their position-matching across boundary and same-context pairs, see **Supplementary Figures 1**-**2**.

Analyses were conducted in RStudio (Version 2023.12.1, Posit Team, 2024), with the main modeling using the lme4 (Bates, Maechler, Bolker, & Walker, 2015) and ordinal packages (Christensen, 2023). Each model included random intercepts for participant ID. Continuous predictors were mean-centered by-participant to reduce the influence of individual differences (see Enders & Tofighi, 2007). This mean centering was performed separately for each model. Continuous predictors were also *z*-scored as necessary to improve model convergence.

## Acknowledgments

This project was funded by federal NIH grant R01 MH074692 to Lila Davachi and by fellowships on federal NIH grant F32 MH114536 to D.C. We thank Jaime Castrellon, Alexandra Cohen, and Jacinda Taggett for very helpful feedback on earlier versions of this manuscript.

## Supplementary Material

**Supplementary Figure 1.**
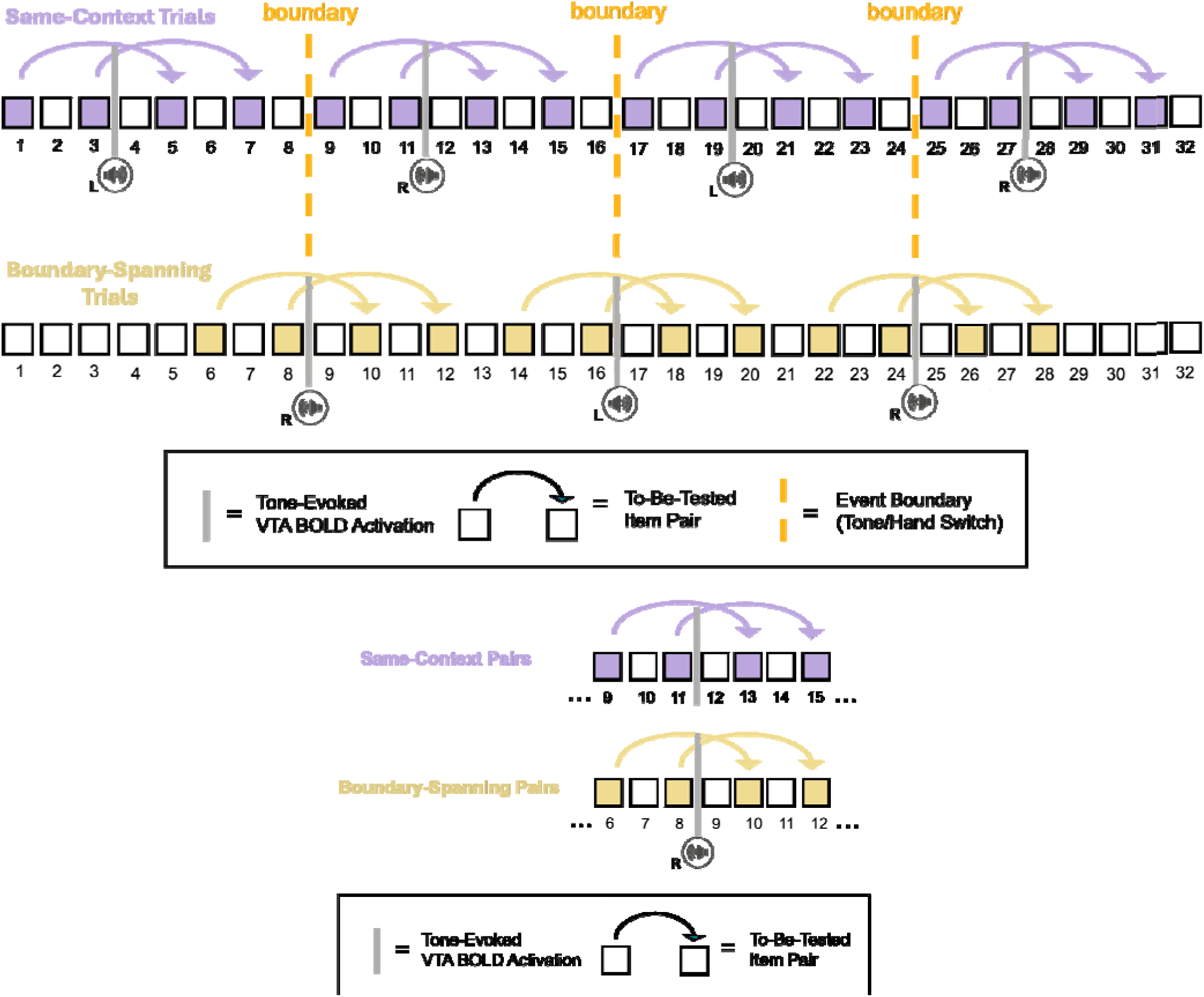
Item pair structure across the event sequence task. (*Top panel*) Each sequence contained 32 images of everyday objects. A pure tone was played either in participants’ left ear or right ear for 8 successive items. It then switched to the other ear and changed in pitch. The side/tone then repeated for another 8 items before switching ears again and so on. Colored squares denote the item pair positions that were subsequently queried during the temporal memory tests after each list. The arrow connections indicate which item pairs were later tested together, in the memory test. There were always three intervening items between each to-be-tested pair during encoding. Purple squares show the pair positions for same-context trials, or to-be-tested item pairs that were presented with the same tone. Yellow squares show the pair positions for boundary trials, or to-be-tested item pairs that spanned an intervening tone switch. Vertical dashed dark yellow lines indicate the positions of “event boundaries”, or the three auditory tone switches in each list. For the linear mixed effects modeling and cumulative link modeling analyses, we aimed to relate trial-level estimates of tone-evoked VTA activation to both temporal distance memory ratings and blinking behavior associated with their corresponding item pairs. To align these two measures appropriately, we specifically focused on VTA parameter estimates (derived from BOLD signal model fits in the GLM) evoked by event boundaries and same-context tones position-matched to those locations (all vertical gray lines). (*Bottom panel*) Example of position-matching between tone-evoked VTA activation between conditions. This illustration shows how the specific tones analyzed for VTA activation were position-matched based on the location of event boundaries. This provided a critical reference point for controlling timing-related effects of VTA activation relative to the positioning of the to-be-tested item pairs spanning those moments. In this example section from an item sequence, the first boundary tone occurred after the 8^th^ item in the list. This means that the boundary occurred after two items for the item from the first pair that spanned those boundaries (i.e., item in position 6), and immediately after the first item in the second boundary-spanning item pair (i.e., item in position 8). To position-match these timing effects in the same-context condition, the two same-context pairs were also temporally aligned with the same sampling points. That is, the relative positioning of the selected tones in the analysis was the same as the two boundary-spanning pairs for each event boundary in the list.

**Supplementary Figure 2.**
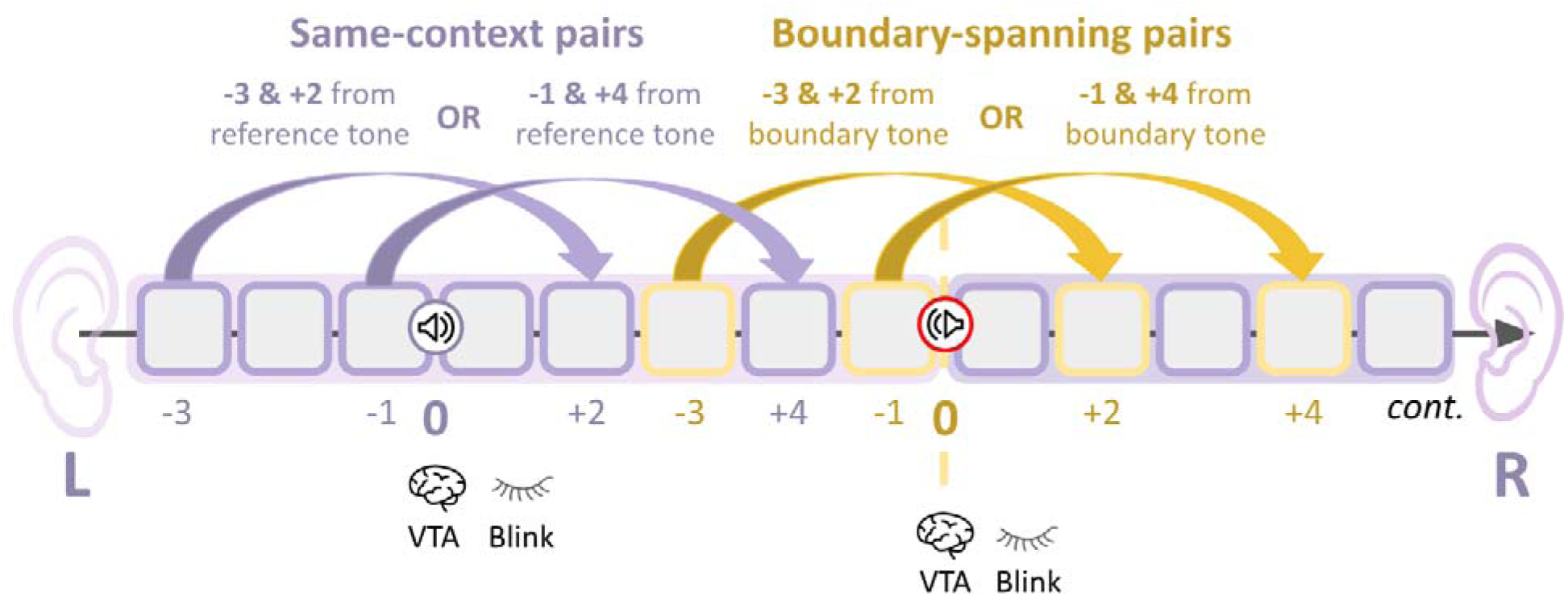
Detailed schematic of position labeling and matching between conditions. Several analyses examined how indirect measures of brief dopaminergic processes during encoding (i.e., tone-related VTA activation or post-tone blink count) predicted larger-scale outcomes (i.e., memory for temporal distance or blink rate between to-be-tested pairs). For these analyses, one intervening tone was selected for each to-be-tested pair in the event encoding sequence. Arrow connections represent to-be-tested pairs, including same-context pairs (purple) and boundary-spanning pairs (yellow). For boundary-spanning pairs, the selected tone was the tone switch that denoted an event boundary (red circle). Depending on the particular boundary-spanning pair, the to-be-tested items fell either 3 positions before (−3) and 2 positions after (+2) the tone switch or 1 position before (−1) and 4 positions after (+4) the tone switch. For same-context pairs, the selected tone (purple circle) matched these positions (i.e., −3 and +2 or −1 and +4).

**Supplementary Figure 3.**
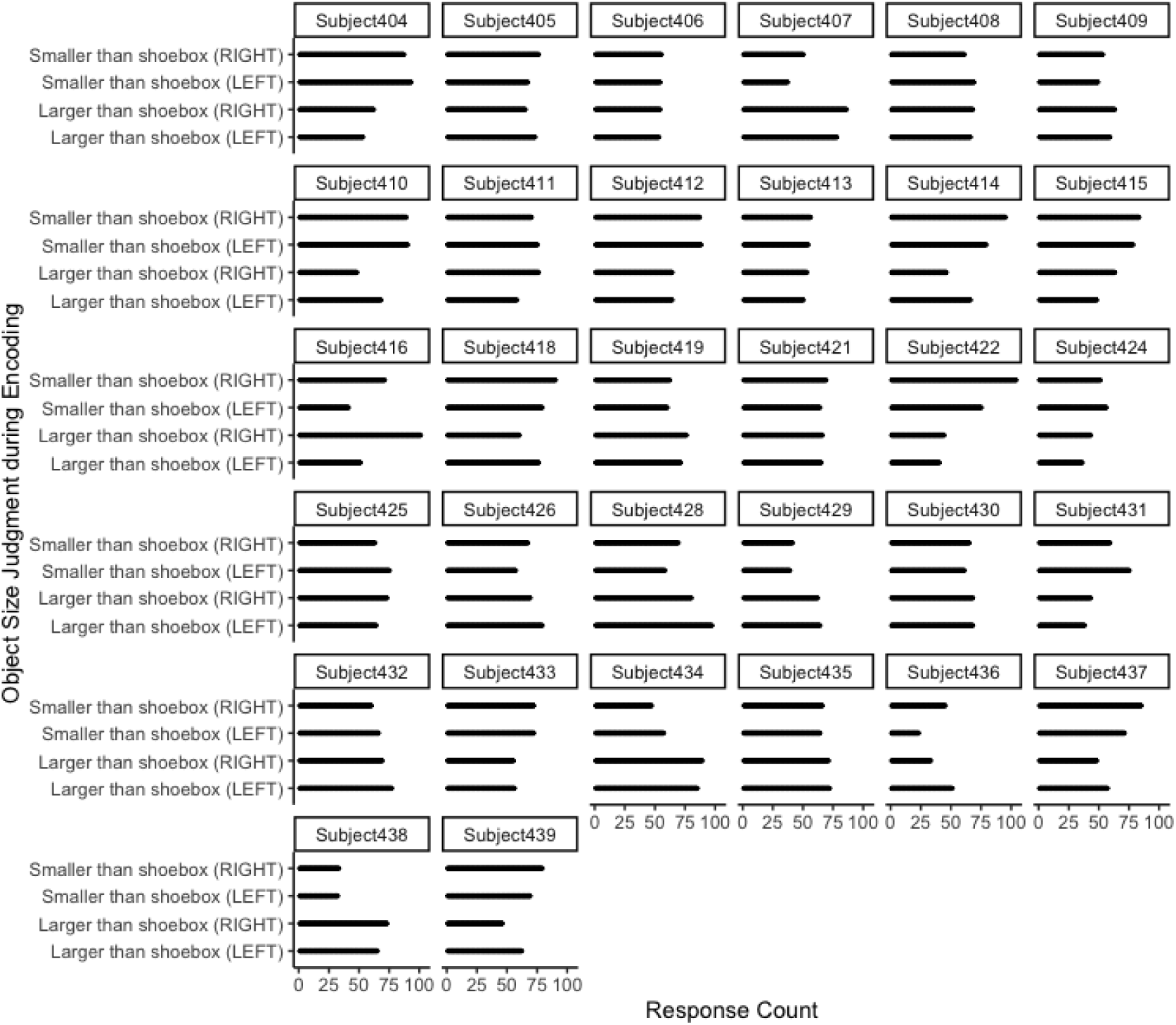
All participants used all four response options during the encoding orienting question. Response distributions for each participant to the simple orienting question: “Is this object larger or smaller than a standard shoebox?”. For each object image, participants were instructed to respond with a button box in their left or right hand, depending on the side of the preceding auditory tone (e.g., right ear = right hand).

**Supplementary Figure 4.**
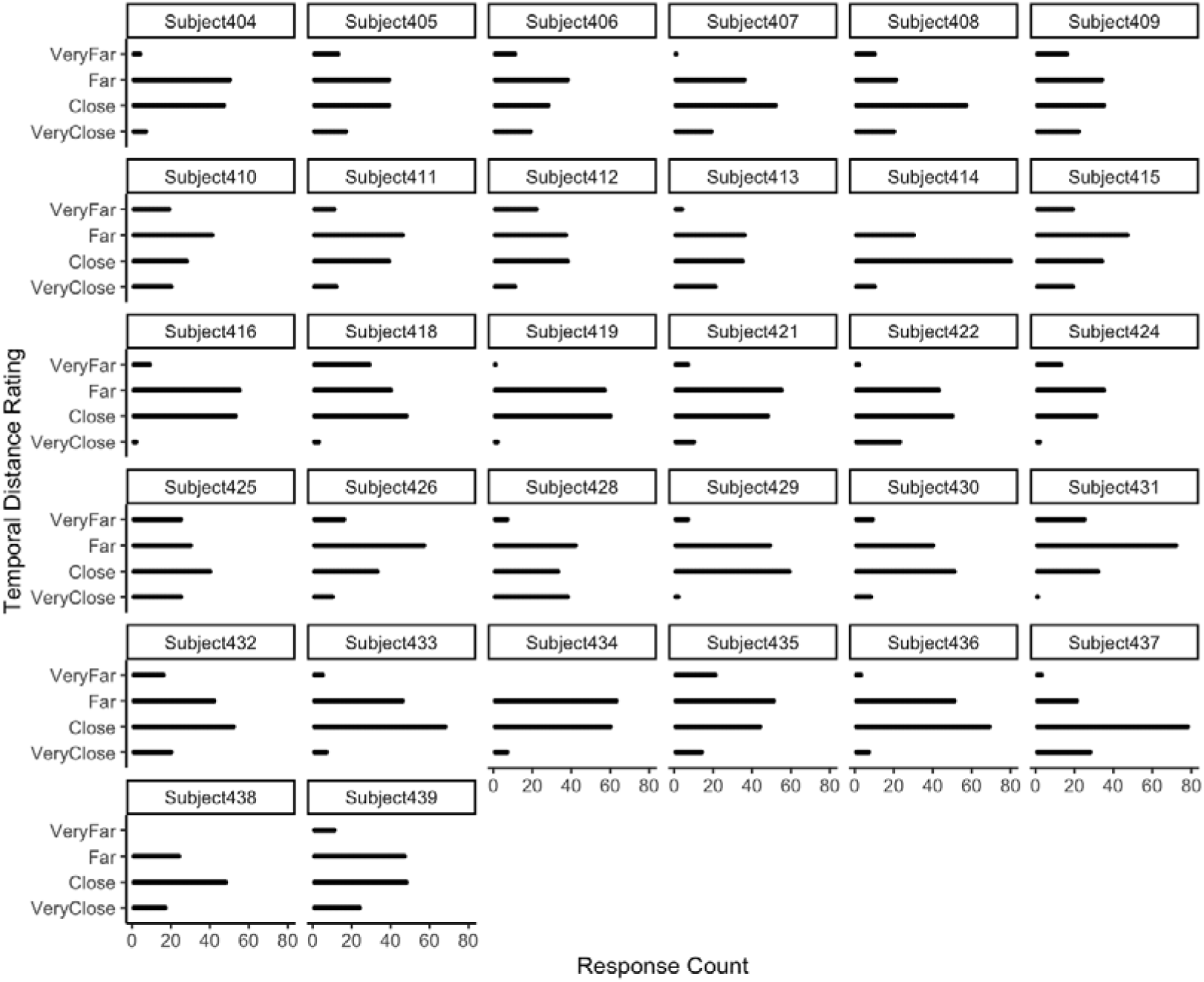
Participants generally used all four rating options on the temporal distance memory test. Response distributions for each participant on the temporal distance memory test. On each test trial, participants were presented with different pairs of objects from the prior sequence and asked to rate how far apart they thought each pair appeared in time (‘very close’, ‘close’, ‘far’, or ‘very far), despite their equivalent objective distance during encoding.

**Supplementary Figure 5.**
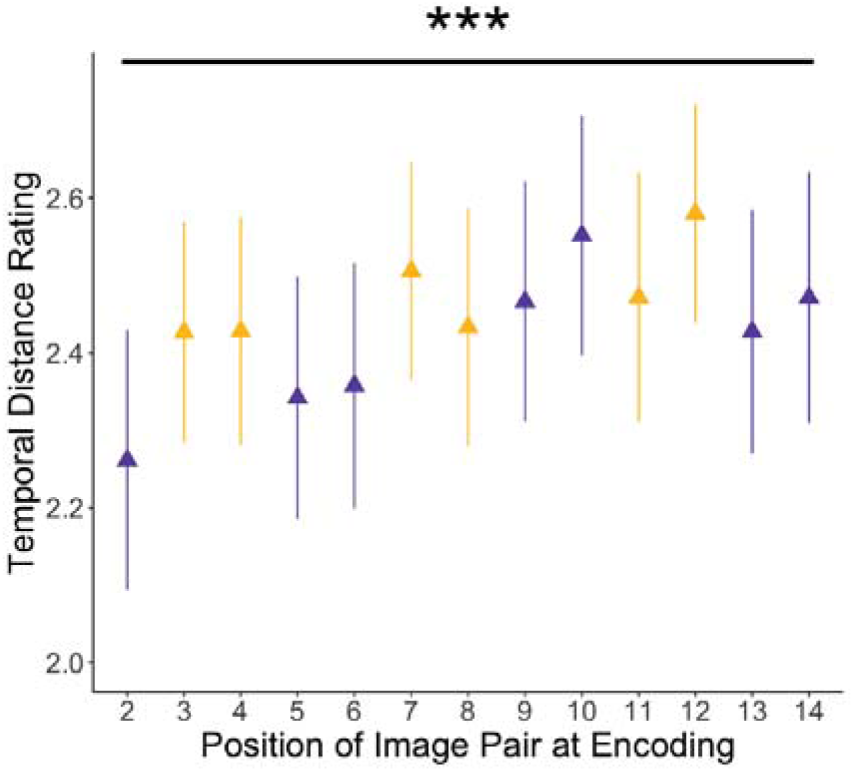
Later temporal distance memory expands over an entire encoding block. Mean ratings on the temporal distance memory test by position of the to-be-tested image pairs across an entire encoding block. Ratings (1-4) are displayed as a continuous variable. Yellow triangles represent boundary-spanning item pairs, while purple triangles represent same-context item pairs. In order of pair presentation at encoding, the boundary-spanning pairs were the 3^rd^, 4^th^, 7^th^, 8^th^, 11^th^, and 12^th^ item pairs. Error bars represent SEM. Statistical significance refers to results from trial-level cumulative link model in which pair position was modeled as an integer fixed-effect predictor of temporal distance ratings (*ß* = .035, *SE* = .0081, *z* = 4.31, *p* < .001). \*\*\**p* < .001.

### Relationships between noradrenergic and dopaminergic system activation measures and other critical temporal distance and eye-tracking results

According to theoretical frameworks of event segmentation (Zacks & Sargent, 2010), both the dopaminergic and noradrenergic systems are well positioned to elicit a global updating signal when event transitions occur. However, it is possible that they contribute to memory separation and blink behavior in different ways (Rouhani et al., 2024). We have also previously shown that LC activation at boundaries relates to impairments in temporal order memory, another index of memory separation, and more differentiated multivoxel activation patterns in left dentate gyrus (DG), a hippocampal subregion that is important for memory encoding and in disambiguating overlapping representations (Clewett et al., 2025). Much work shows that dopaminergic modulation of hippocampal processing also plays an essential role in structuring and encoding episodic memories, making it another candidate input signal for memory separation at its target regions (Shohamy & Adcock, 2010). In the next set of exploratory analyses, we sought to determine potential differences and similarities between these systems’ contributions to temporal memory and links to different eye-tracking measures.

### Locus coeruleus (LC) region-of-interest (ROI) definition

The anatomy of the LC was visualized in each participant using a fast spin echo (FSE) T1 MRI sequence, which is sensitive to neuromelanin and water content within LC neurons. Each participant’s left and right LC were hand-drawn separately by trained raters and then transformed into each participant’s run-level functional space to extract parameter estimates of trial-level LC activation. For more details about this MRI sequence, drawing procedures, and analysis methods, see Clewett et al. (2025).

### Testing the relationship between tone-related LC activation and temporal distance ratings

To assess whether temporal distance memory could also be predicted by LC activation, we fit a cumulative link model where tone-related LC activation was mean-centered by participant and modeled as a fixed-effect predictor. Condition was modeled as a categorical variable, and we also included an interaction term between the two. Pair position at encoding was modeled as an integer fixed-effect predictor. Participant ID was modeled as a random effect. The results revealed no significant main effect of LC activation on temporal distance ratings (*ß* = .018, *SE* = .031, *z* = .59, *p* = .56) nor significant LC-by-pair type interaction effect (*ß* = .018, *SE* = .031, *z* = .58, *p* = .56) (see **Supplementary Figure 6** below for results for each pair type separately).

### Comparing relationships between tone-related LC vs. VTA activation and distance memory

Next, we clarified whether the coupling between brainstem activation and temporal distance memory for boundary pairs was specific to the VTA. We performed a likelihood-ratio test that compared a full model with both VTA and LC activation as predictors with a restricted model that averaged VTA and LC activation as one predictor. The test revealed that the full model did not significantly improve fit over the restricted model (χ^2^ (1) = .83, *p* = .36). Therefore, we cannot conclude that the relationship with temporal distance for boundary pairs is unique to VTA activation.

**Supplementary Figure 6.**
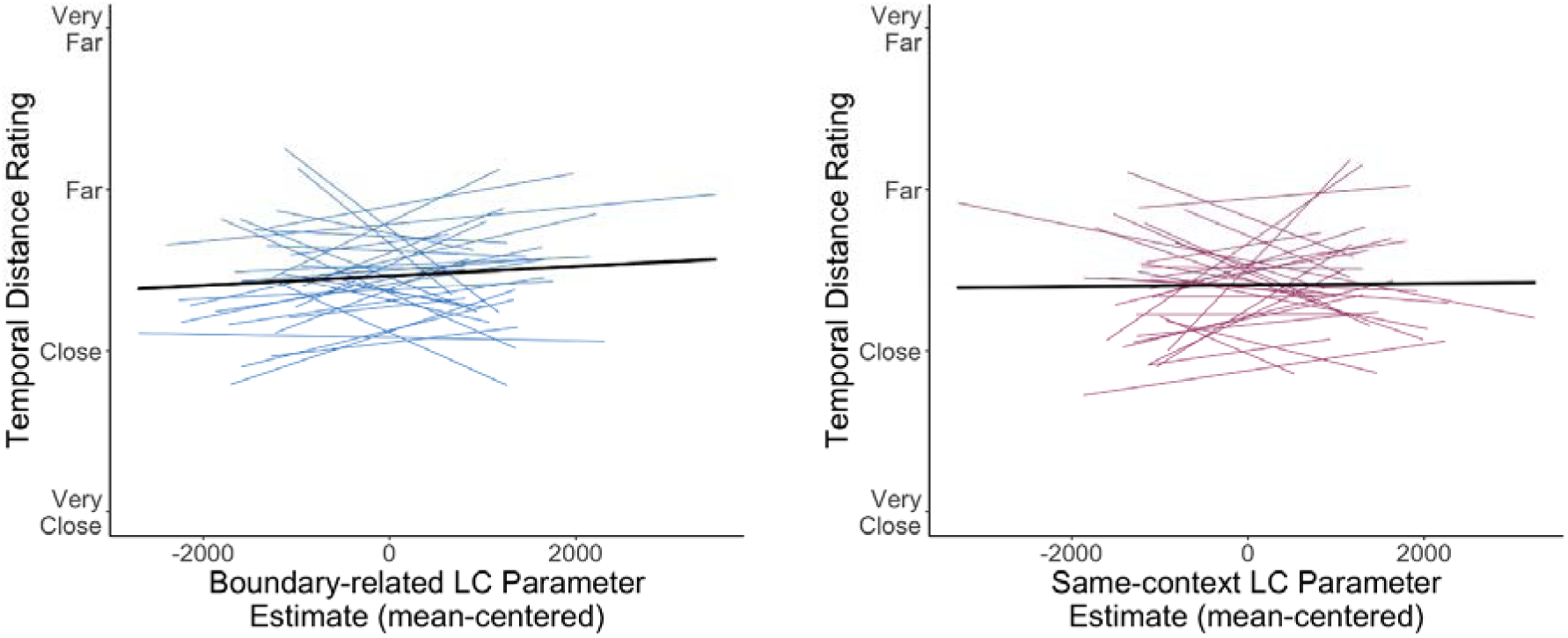
Tone-related LC activation did not predict subjective temporal distance memory between item pairs. (*Left panel*) Participant-level trendlines plotting the association between boundary-related LC parameter estimates (mean-centered) and temporal distance memory ratings (displayed as a continuous variable). (*Right panel*) Participant-level trendlines plotting the association between same-context LC parameter estimates and temporal distance memory ratings. Dark, bold lines represent the average linear trend across participants.

### Testing the relationship between tone-related VTA activation and temporal order memory

To determine whether temporal order accuracy could also be predicted by VTA activation, we fit a generalized linear mixed effects model that included tone-related VTA activation as a fixed-effect predictor of order accuracy (correct = 1, incorrect = 0). Pair position at encoding was modeled as an integer fixed-effect predictor. We found no significant main effect of VTA activation (*ß* = .037, *SE* = .035, *z* = 1.05, *p* = .30) or VTA-by-pair type interaction effect (*ß* = .014, *SE* = .035, *z* = .40, *p* = .69) on order accuracy (**Supplementary Figure 7**).

### Comparing relationships between tone-related LC vs. VTA activation and order memory

Next, we clarified whether the coupling between brainstem activation and temporal order ratings for boundary pairs was specific to the LC. We performed a likelihood-ratio test that compared a full model with both VTA and LC activation as predictors with a restricted model that averaged LC and VTA activation as one predictor. To avoid a singular fit, these models were fit with a fixed effect representing the side of the screen containing the correct answer (i.e., left or right). The test revealed that the full model significantly improved fit over the restricted model (^2^(1) = 9.21, *p* = .0024). This finding suggests that the relationship between brain activation and temporal order memory for boundary-spanning pairs was specific to the LC.

### Comparing relationships between temporal order vs. distance memory and VTA activation

T compare the magnitude of the relationship between tone-related VTA activation and temporal memory for boundary pairs, we compared the standardized coefficients for VTA activation between the distance and order memory models. To avoid a singular fit, the order memory model was fit with a fixed effect representing the side of the screen containing the correct answer (i.e., left or right). The standardized coefficient for VTA activation in the distance memory model (*ß*\* = .10, 95% CI: [.01, .20]) was not significantly different than in the order memory model (*ß*\* = .05, 95% CI: [-.05, .16]). Thus, while VTA activation did not relate to temporal order memory, we cannot conclude that VTA coupling with memory was unique to temporal distance memory.

**Supplementary Figure 7.**
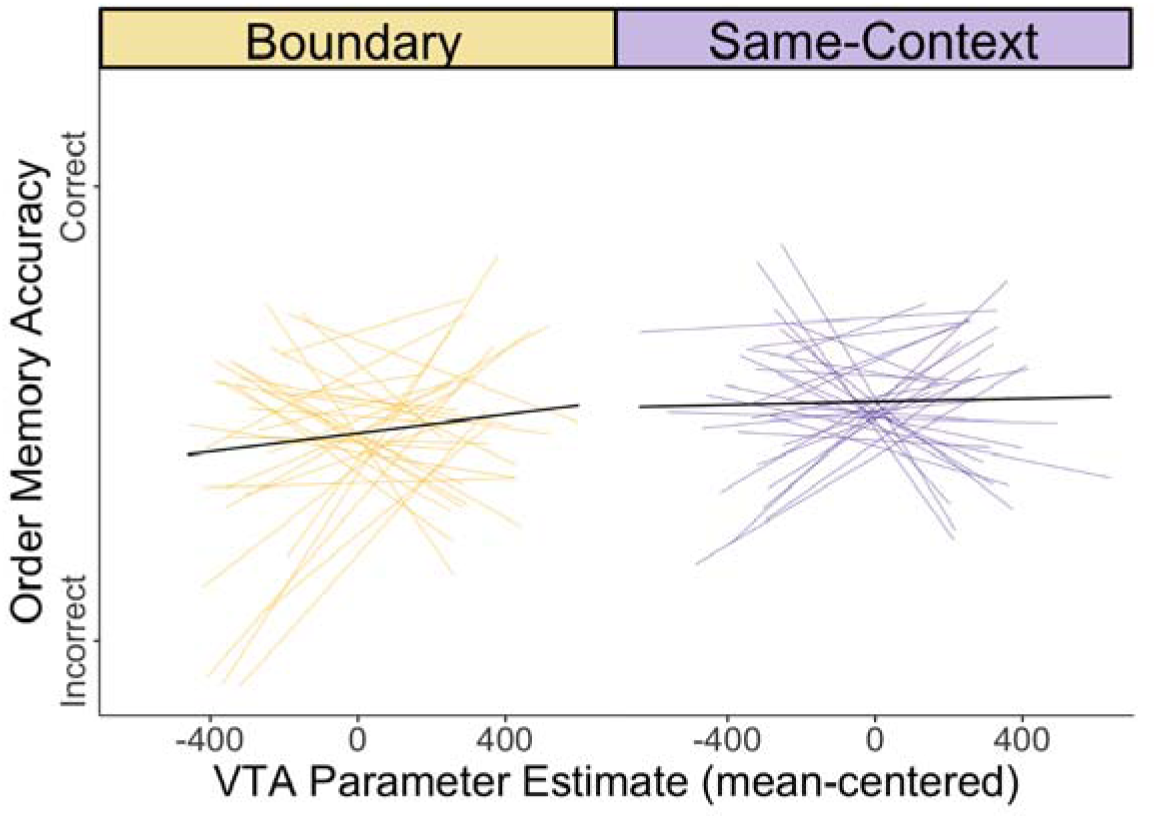
Tone-related VTA activation did not predict temporal order memory accuracy. Participant-level trendlines plotting the null relationship between VTA parameter estimates (mean-centered) and order memory accuracy for boundary-spanning pairs (yellow; left) and same-context pairs (purple; right). For modeling, VTA parameter estimates were mean-centered for each condition separately. Order memory accuracy is displayed as a continuous variable. Dark, bold lines represent the average linear trend across participants.

### Testing the relationship between blinking and temporal order memory

To determine whether temporal order accuracy could also be predicted by blinking, we fit a generalized linear mixed effects model that included extended blink count as a fixed-effect predictor of order accuracy (correct = 1, incorrect = 0). We found no significant main effect of extended blink count (*ß* = -.0024, *SE* = .0044, *z* = - .55, *p* = .58) or blink count-by-pair type interaction effect (*ß* = -.0052, *SE* = .0044, *z* = −1.18, *p* = .24) on order accuracy (**Supplementary Figure 8**).

### Comparing relationships between temporal order vs. distance memory and blinking

To compare the magnitude of the relationship between blinking and temporal memory ratings for boundary pairs, we compared the standardized coefficients for blinking between the distance and order memory models. To avoid a singular fit, the order memory model was fit with a fixed effect representing the side of the screen containing the correct answer (i.e., left or right). The standardized coefficient for blinking in the distance memory model (*ß*\* = .16, 95% CI: [.06, .27]) was significantly greater than in the order memory model (*ß*\* = -.07, 95% CI: [-.18, .04]). This finding suggests that the relationship between blinking and memory for boundary-spanning pairs was greater for temporal distance memory than order memory.

**Supplementary Figure 8.**
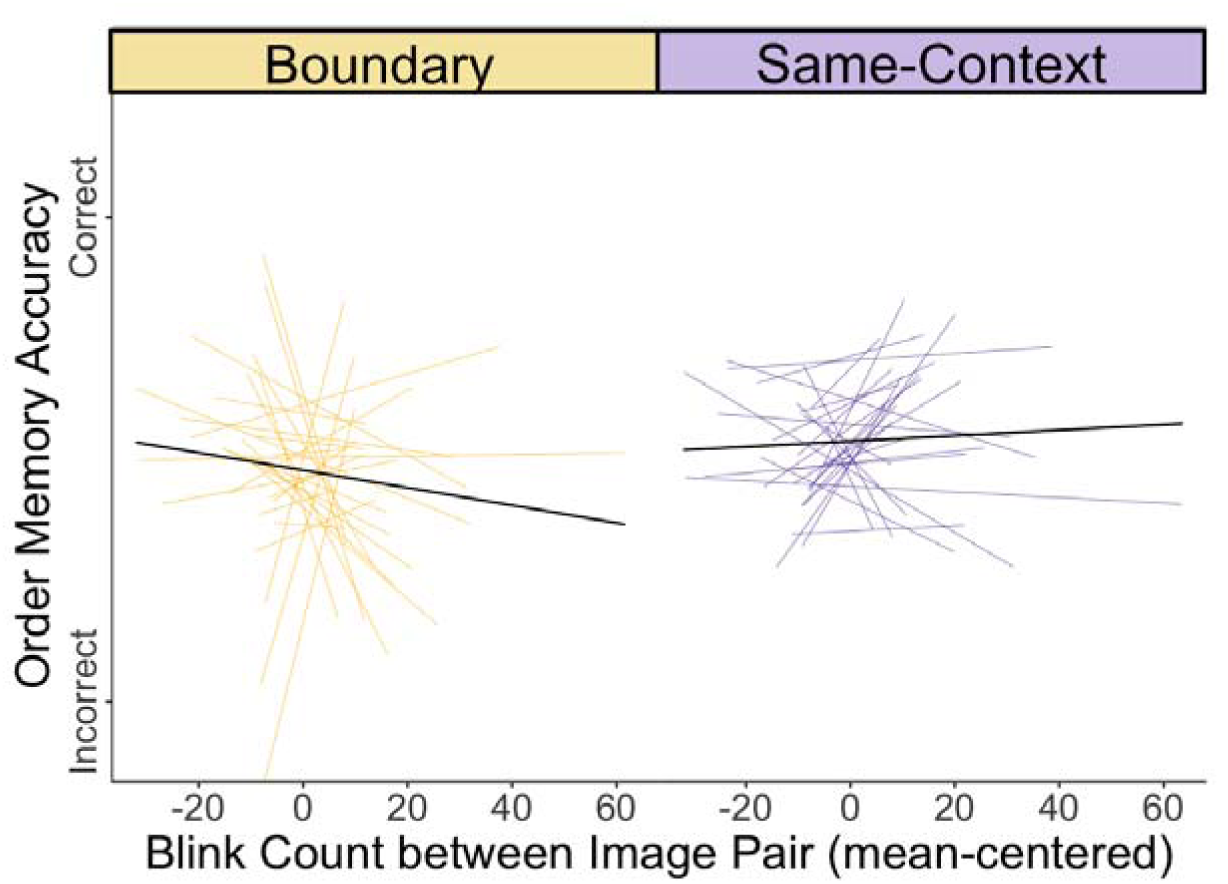
Sustained blinking between to-be-tested item pairs did not predict memory accuracy for the temporal order of those items. Participant-level trendlines plotting the null relationship between extended blink count (mean-centered) and order memory accuracy for boundary-spanning pairs (yellow; left) and same-context pairs (purple; right). For modeling, blink counts were mean-centered for each condition separately. Order memory accuracy is displayed as a continuous variable. Dark, bold lines represent the average linear trend across participants.

### Testing the relationship between tone-related LC activation and blinking

To assess whether blinking behavior could also be predicted by LC activation, we fit linear mixed effects models. First, we examined local blink count. We found no significant main effect of LC activation on post-tone blink count (*ß* = -.026, *SE* = .018, *t*(7308.17) = −1.44, *p* = .15; **Supplementary Figure 9, left panel**). There was also no significant LC-by-pair type interaction effect on blinking (*ß* = -.018, *SE* = .018, *t*(7308.32) = -.97, *p* = .33).

Next, we examined temporally extended blink count between to-be-tested image pairs. We found no significant main effect of LC activation on extended blink count (*ß* = .00025, *SE* = .00025, *t*(2806) = 1.00, *p* = .32; **Supplementary Figure 9, right panel**). There was also no significant LC-by-pair type interaction effect (*ß* = .00023, *SE* = .00025, *t*(2806) = .90, *p* = .37).

### Comparing relationships between tone-related VTA vs. LC activation and blinking

Next, we clarified whether the coupling between brainstem activation and blinking between to-be-tested pairs was specific to the VTA. We performed a linear hypothesis test that compared a full model with both VTA and LC activation as predictors with a restricted model in which the VTA and LC coefficients were equal. The test revealed that these coefficients were not equal (^2^(1) = 6.95, *p* = .0084), suggesting that the relationship between brain activation and temporally extended blink count was specific to the VTA.

**Supplementary Figure 9.**
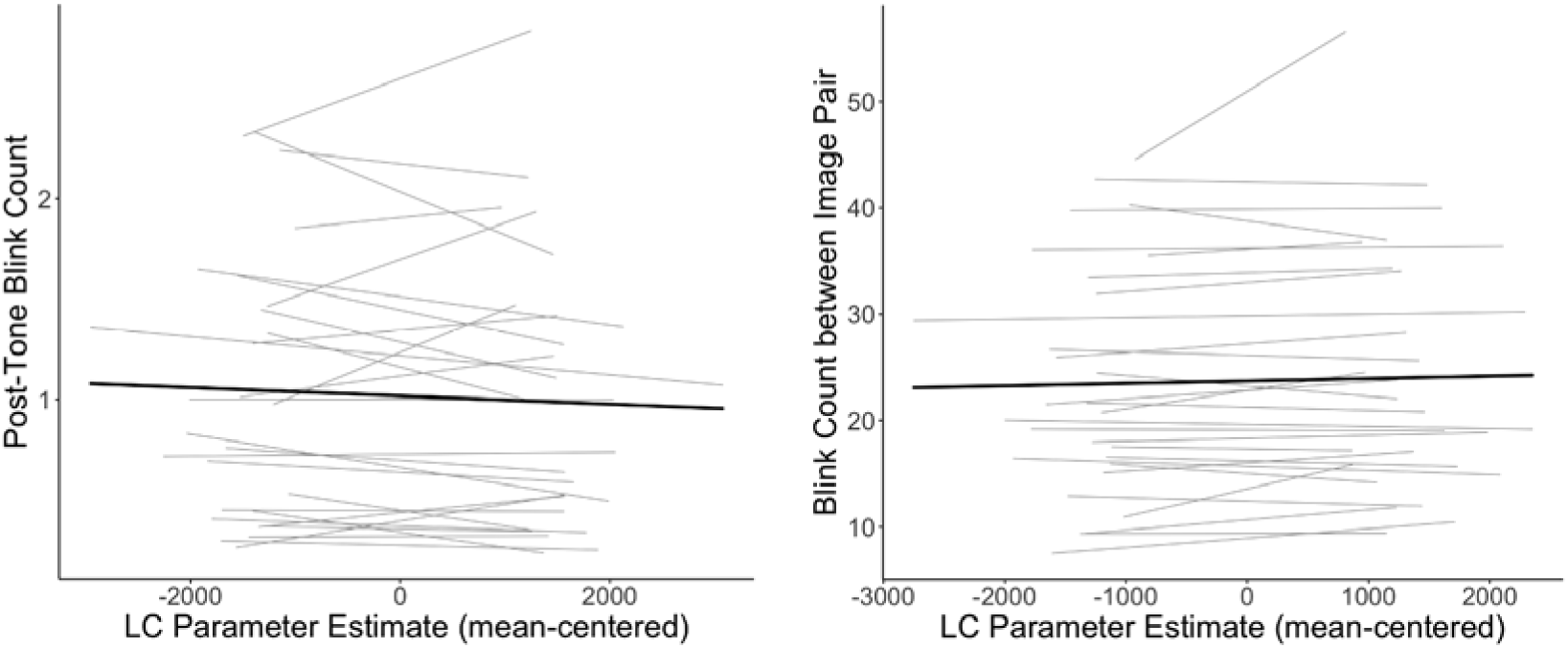
Tone-related LC activation did not predict momentary or temporally extended changes in blink behavior. (*Left panel*) Participant-level trendlines plotting the null relationship between LC parameter estimates and post-tone blink count. (*Right panel*) Participant-level trendlines plotting the null relationship between LC parameter estimates and extended blink count. Dark, bold lines represent the average linear trend across participants. LC parameter estimates were mean-centered for each model separately.

### Hippocampal pattern similarity fMRI analyses

Left and right hippocampal subfields CA2/3, dentate gyrus (DG), and CA1 were segmented from each participant’s high-resolution anatomical scan using Freesurfer 6.0 (https://surfer.nmr.mgh.harvard.edu/). Validated hippocampal ROIs were then co-registered to each participant’s native/run-specific functional space and thresholded at 0.2 to reduce spatial overlap between adjacent subfields.

For each of these hippocampal ROIs, we extracted activation patterns from the trial-unique beta maps produced by the LSS GLM, which modeled stimulus-specific activation patterns for all tones and images in a sequence. Here, we focused on the multivoxel patterns evoked by the image pairs as an index of hippocampal pattern stability across encoding, with more similar patterns reflecting representational stability and more dissimilar patterns reflecting temporal pattern separation. Hippocampal subfield pattern similarity scores were computed at the item pair level by correlating multivoxel patterns between each of the to-be-tested trial pairs from encoding. For more details, see Clewett et al. (2025).

### Testing the relationship between boundary-related VTA activation and hippocampal pattern similarity across time

To determine whether VTA activation at boundaries predicted hippocampal pattern similarity for to-be-tested item pairs spanning those same boundaries, we fit a linear mixed effects model with six predictors of VTA activation at boundaries: left and right DG, CA1, and CA2/3 pattern similarity. Unlike the LC (Clewett et al., 2025), left DG pattern similarity did not significantly predict tone-induced VTA activation (*ß* = −6.70, *SE* = 51.44, *t*(1348.17) = -.13, *p* = .90). There were also no significant main effects of right DG, left or right CA1, or left or right CA2/3 (*p*s > .05) (all subfield results are displayed in **Supplementary Figure 10**).

### Comparing relationships between boundary-related LC vs. VTA activation and hippocampal pattern similarity

To directly compare the effect of left DG pattern similarity on boundary-related LC and VTA activation, we added brain region (LC vs. VTA) as an interaction term in the model. We observed a significant region-by-left DG interaction effect (*ß* = −414.35, *SE* = 120.49, *t*(2732) = −3.44, *p* < .001). A simple slopes analysis revealed that the slopes of left DG pattern similarity significantly differed by brain region (*t* ratio = −3.44, p < .001), such that LC activation at boundaries was more strongly linked to left DG pattern similarity (*b* = −841.9) than VTA activation (*b* = −13.2). There were no other significant interaction effects (*p*s > .05).

In summary, these findings suggest that momentary increases in VTA activation at boundaries did not modulate pattern similarity in hippocampal subfields. We also found that LC activation (vs. VTA) at boundaries was more strongly coupled with left DG pattern similarity for items spanning those boundaries, suggesting the LC may play a larger role in differentiating memory representations across time. Moreover, LC activation was selectively coupled with impairments in temporal order memory across boundaries, suggesting that the noradrenergic system might specifically modulate objective features of temporal memory.

**Supplementary Figure 10.**
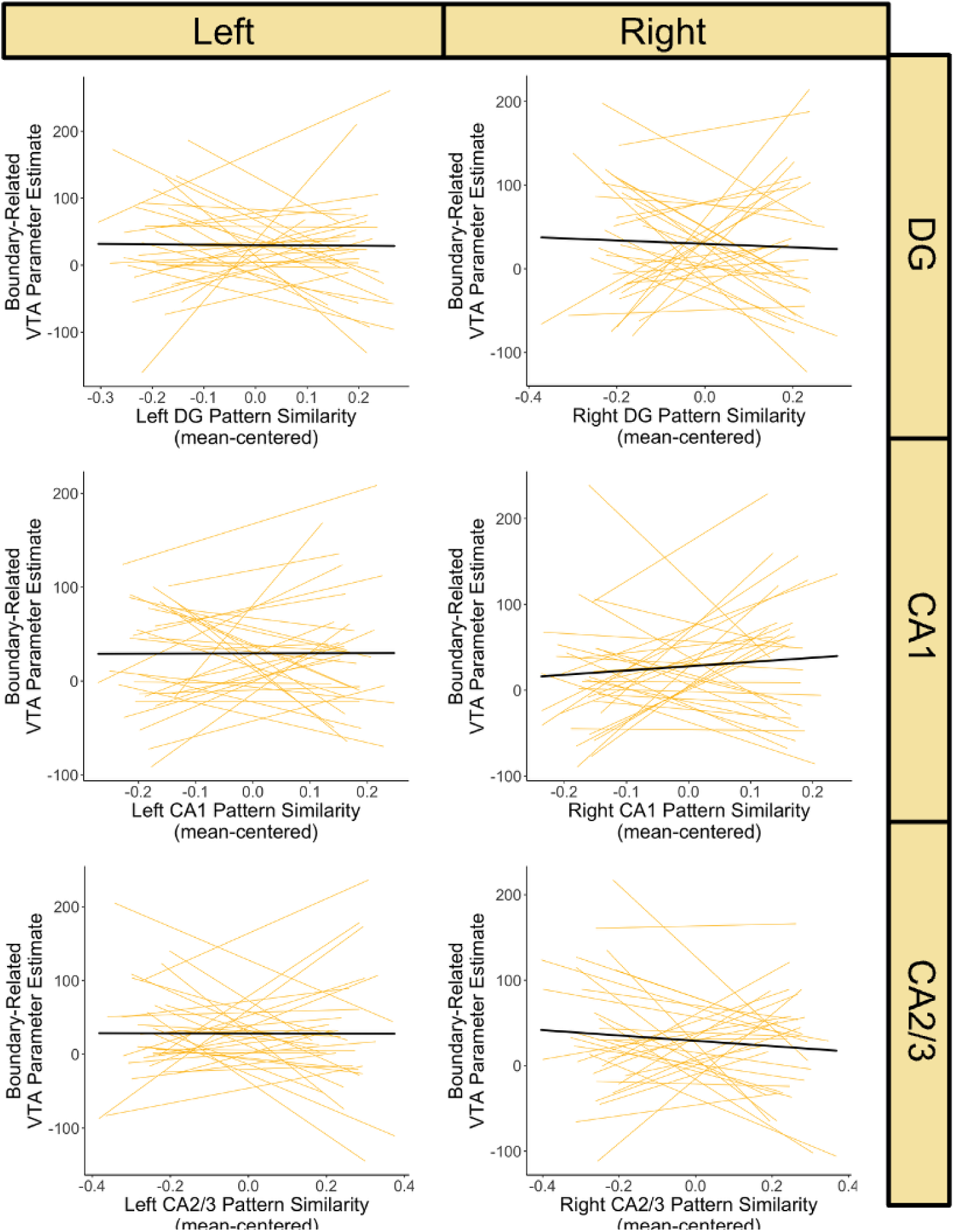
Hippocampal pattern similarity for item pairs, including in the dentate gyrus, does not predict boundary-related VTA activation. Participant-level trendlines plotting the null relationships between hippocampal subfield pattern similarity for to-be-tested item pairs (in dentate gyrus, or DG, CA1, and CA2/3; left and right) and boundary-related VTA parameter estimates between those same item pairs. Dark, bold lines represent the average linear trend across participants. Each subfield’s pattern similarity values were mean centered separately and were all included as predictors of boundary-related VTA activation in the same linear mixed effects model.

### Pupil dilation temporal principal component analysis (PCA)

Prior work shows that context shifts elicit increased pupil dilation, suggesting that boundaries engage central arousal processes (Clewett et al., 2025). Pupil dilation, however, is complex and is mediated by multiple autonomic pathways and neuromodulatory systems (Reimer et al., 2016). Building on earlier work (Clewett et al., 2020), we use a temporal principal component analysis (PCA) to decompose boundary-related pupil dilations into its distinct temporal features, providing a unique opportunity to link event segmentation to different neural and behavioral effects. The temporal PCA on tone-evoked pupil dilations revealed three canonical pupil components identified in prior work, including a biphasic response that may index separate influences of parasympathetic and sympathetic nervous system regulation on pupil diameter (Clewett et al., 2020; Steinhauer and Hakerem, 1992). The temporal characteristics of these pupil components, including their latencies-to-peak and percent of explained variance, were as follows: (1) an early-peaking component (684 ms; 89.26% variance); (2) intermediate-peaking component (1,420 ms; 8.40% variance); and (3) slowly decreasing component (19.6 ms; 1.27% variance). For more details about these methods and results, see Clewett et al. (2025).

### Testing the relationship between boundary-related VTA activation and three distinct temporal features of tone-evoked pupil dilation

In previously published work using this dataset, we found that pupil components #2 and #3 were both positively coupled with boundary-related LC activation (Clewett et al., 2025). Here, we tested whether the pupil components were also correlated with engagement of the VTA. Using Spearman’s rho correlations, we found that boundary-induced VTA activation was not significantly correlated with boundary-induced loading on pupil component #1 (III = .14, *p* = .47), pupil component #2 (III = .11, *p* = .58), or component #3 (III = -.079, *p* = .69; **Supplementary Figure 11**).

### Comparing the relationships between boundary-related VTA vs. LC activation and pupil dilation

Next, we examined whether the coupling between brainstem activation and the three pupil components was specific to the LC using a Steiger’s Z test. We found no significant differences in the linear relationships between LC and VTA activation and pupil component #1 (*z* = -.44, *p* = .66), pupil component #2 (*z* = -.44, p = .66), or pupil component #3 (*z* = .48, *p* = .63). Thus, while VTA activation did not relate to distinct temporal characteristics of pupil dilation at boundaries, we cannot conclude that neuromodulatory coupling with pupil responses was unique to LC activation.

**Supplementary Figure 11.**
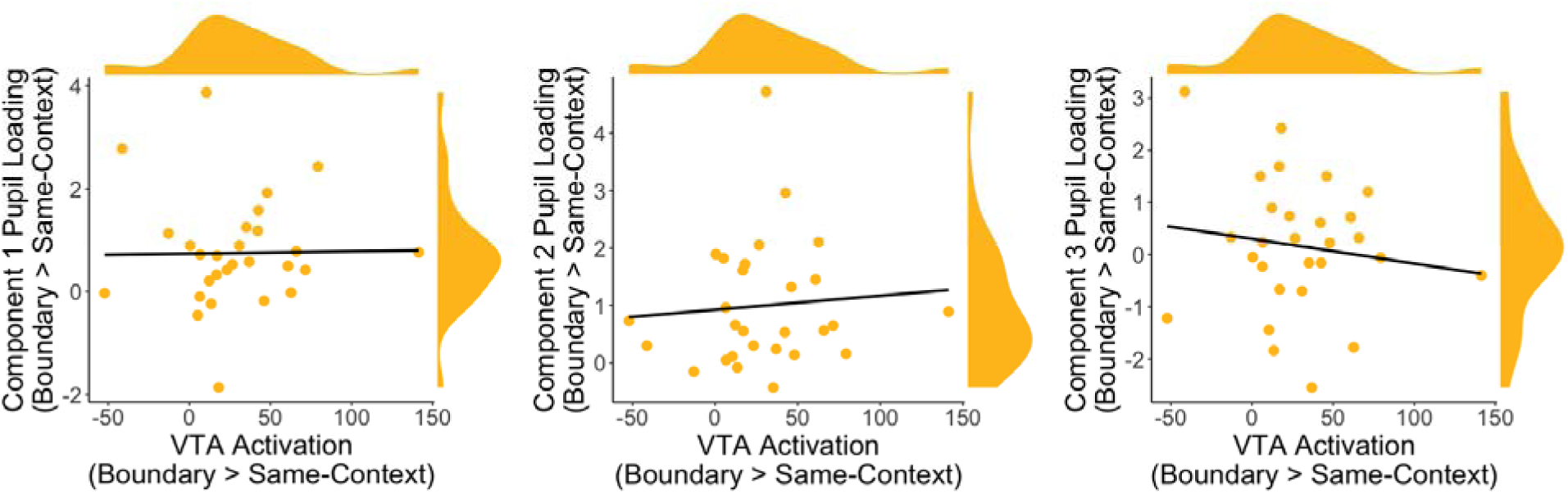
Boundary-related VTA activation was not associated with any of three temporal features of tone-evoked pupil dilations. Spearman’s rho correlation plots showing that boundary-induced VTA activation was not significantly correlated with boundary-related engagement of pupil dilation component #1 (*left panel*), component #2 (*middle panel*), or component #3 (*right panel*). Individual dots represent each participants’ data. *X*- and *y*-distributions also displayed.

### Identifying which blinking periods and types of stimulus-elicited blinked predicted time dilation in memory

In this set of analyses, we aimed to identify the specific blink periods and stimuli that predict later distortions in temporal distance memory.

### Average patterns of blinking across the inter-pair windows

To capture temporally dynamic patterns of blinking across the event sequence, we first divided each inter-item window into two coarse-grained intervals: 1) the interval *before* the reference tone (*M* duration = 12.28s); and 2) the interval *after* the reference tone (including that tone; *M* duration = 20.87s). For more details, see **Supplementary Figure 12** below. On average, participants blinked 8.83 times before the reference tone (*M* boundary pairs = 8.78 blinks; *M* same-context pairs = 8.87 blinks) and 15.43 times after the reference tone (*M* boundary pairs = 15.77 blinks; *M* same-context pairs = 15.10 blinks).

Next, given that the interval *after* the reference tone carried key contextual information in this task (e.g., that the auditory-task context had or had not changed), we divided this interval further. Specifically, we identified even more fine-grained intervals associated with each stimulus: 1) images; and 2) tones (**Supplementary Figure 12**). We observed that most blinks occurred after images (*M* overall = 12.56 blinks across images; *M* boundary pairs = 12.77 blinks; *M* = same-context pairs = 12.34 blinks) compared to tones (*M* overall = 4.65 blinks across tones; *M* boundary pairs = 4.76 blinks; *M* = same-context pairs = 4.54 blinks). This is a sensible result, given that images have a longer duration and subsequent ISI compared to tones.

### Relating specific blink periods to temporal distance memory

Here, we conducted linear mixed-effects models to test which stimulus-evoked blinks and periods of blinking were significantly predictive of temporal distance ratings.

### Blinks in the earlier vs. later interval did not influence temporal memory predictions

First, we tested whether the interval before versus after the reference tone drove this effect. There was no significant interaction between blink count, Interval Type (before vs. after reference tone), and Pair Type (boundary vs. same-context) on distance memory (*ß* = .0015, *SE* = .0032, *z* = .45, *p* = .65).

### Focusing on the later interval, blinking after a tone switch vs. no switch predicted greater time dilation in memory

While the prior interaction effects were null, we next focused specifically on the blink window *after* the reference tone, because this carried the critical information about the auditory-task context, In this period, we found a significant interaction between blink count and Pair Type on distance memory (*ß* = .0085, *SE* = .0040, *z* = 2.12, *p* = .034). To break down this interaction effect, we then examined the two Pair Types separately. For boundary pairs, there was a marginally significant main effect of post-tone switch blink count on distance memory (*ß* = .011, *SE* = .0060, *z* = 1.87, *p* = .061), such that more blinking after the tone switch predicted greater subsequent time dilation between the to-be-tested object pairs. In contrast, for same-context pairs, there was no significant effect of post-reference tone blink count on distance memory (*ß* = -.0057, *SE* = .0055, *z* = −1.04, *p* = .30). Therefore, blinks after the reference tone predicted later time dilation only when there had been a tone switch, denoting a new auditory context.

### Investigating whether image- or tone-evoked blink effects in the later inter-pair interval predicted time dilation in memory

Zooming in on this later interval further, we asked whether stimulus type mattered; that is, whether blinks following either *images* or *tones* were the driving factor behind changes in temporal distance ratings. We found that there were no interaction effects between blink count, Stimulus Type (Images vs. Tones), and Pair Type on distance memory (all *p*s > .05). Thus, at least during the critical latter part of the inter-pair interval, blinks following images vs. tones did not relate to temporal distance memory.

Given the lack of an interaction effect and that tones carried the important contextual information during the task, we next focused on tone-related blinking alone in the later inter-pair interval. We found a significant interaction between Pair Type and total blink count following tones on distance memory (*ß* = .031, *SE* = .012, *z* = 2.63, *p* = 0.0086). To break down this interaction effect, we analyzed the two Pair Types separately. For boundary pairs, there was a significant main effect of total blink count following tones on distance memory (*ß* = .036, *SE* = .017, *z* = 2.11, *p* = .035), such that more blinking after the tone switch predicted greater time dilation between the to-be-tested object pairs. In contrast, for same-context pairs, there was no significant effect of total blink count following tones on distance memory (*ß* = -.025, *SE* = .016, *z* = −1.56, *p* = 0.12). Together, these findings suggest that the coupling between blinking and time dilation in memory was driven by tone switches, the stimulus that carried the critical signal denoting an event boundary.

In summary, our analyses showed that blinks following a tone switch (versus no switch) predicted greater time dilation in memory between the to-be-tested object pairs. This link between blinking and memory distortion was related to the occurrence of tones, suggesting that event boundaries play an important role in triggering dopaminergic processes that shape later memory separation effects.

**Supplementary Figure 12.**
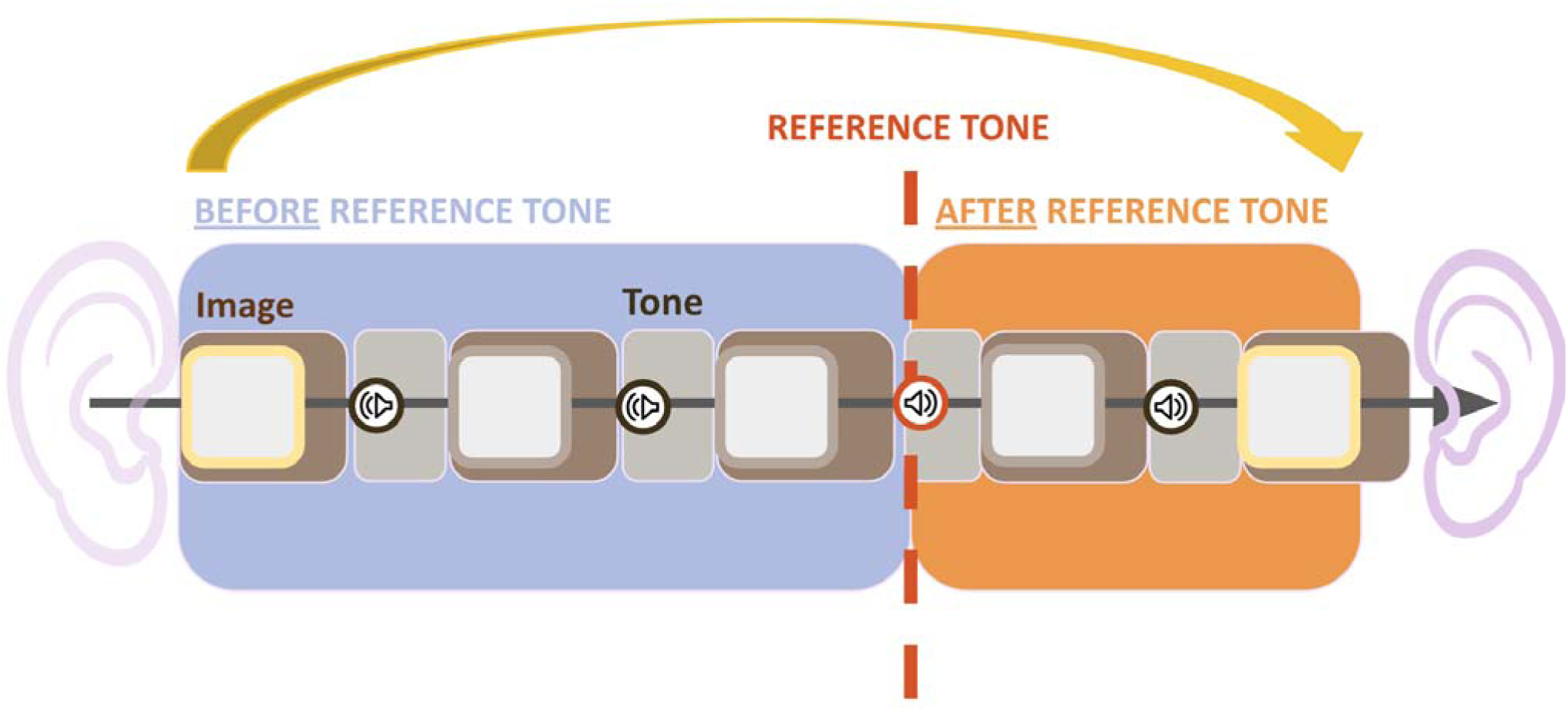
Dividing the window between to-be-tested pairs into specific intervals of interest. Example intervening window for each to-be-tested image pair (see yellow arrow connecting pair of squares). Each window was approximately 32.5s long, containing 5 images and 4 tones total. These windows can be divided into coarse- and fine-grained intervals of interest. The coarse-grained intervals are separated by the reference tone (red dashed line), which is the tone of interest between each to-be-tested pair. For boundary-spanning pairs (example shown here), this was the tone switch that denoted an event boundary. For same-context pairs, this was a position-matched tone (for more details, see **Supplementary** Figure 1**, bottom panel**). Therefore, the two resulting intervals are as follows: (1) *Before* the reference tone (blue), from the onset of the first to-be-tested image to the onset of the reference tone; and (2) *After* the reference tone (orange), from the onset of the reference ton to the offset of the second to-be-tested image. Additionally, the window can be segmented further into fine-grained intervals that are associated with each individual stimulus: (1) Image intervals (brown), from the onset of the image to the end of the subsequent ISI (2.5s + variable + 0.5s); and (2) Tone intervals (gray), from the onset of the tone to the end of the subsequent ISI (variable). The final image interval (brown) extends partially outside of the “*After* the reference tone” (orange) interval, as this image interval also contains the ISI after the final image offset.

